# Suspension culture drives metabolic rewiring in human pluripotent stem cells

**DOI:** 10.64898/2026.09.24.753129

**Authors:** Astrid Sophie Pejstrup Agger, Eleni Kafkia, Mads Bjerknæs Larsen, Thomas Moritz, Christian Toft Madsen, Kristian Honnens de Lichtenberg, Caroline Halloin, Palle Serup, Jan Jakub Zylicz, Christian Honoré

## Abstract

Human pluripotent stem cells (hPSCs) are increasingly expanded in 3D suspension culture to support scalable cell culturing, yet how the transition from 2D to 3D cultures affects hPSCs remains poorly understood. Here, we compare hPSCs in 2D and 3D cultures using transcriptomics, proteomics, spent-medium analysis, and stable-isotope tracing. hPSCs maintained pluripotency in both formats, although suspension culture showed altered growth dynamics and increased initial cell loss. Multi-omics analysis identified culture format as the dominant driver of molecular variation and revealed changes in adhesion, cytoskeletal organization, glycolysis, and oxidative phosphorylation. Spent-medium analysis showed increased lactate/glucose ratio in 3D suspension. Stable-isotope tracing revealed increased glucose-derived labelling of pyruvate and altered partitioning of glucose-derived carbon within the TCA cycle in 3D suspension cultures. Together with increased abundance of glycolytic enzymes, these findings support increased glycolysis in 3D suspension. Glutamine contribution to the TCA cycle was reduced, whereas relative reductive carboxylation was increased in 3D suspension cultures. Together, these findings show that transition from 2D to 3D suspension culture induces metabolic rewiring in hPSCs.

**Graphical abstract:** 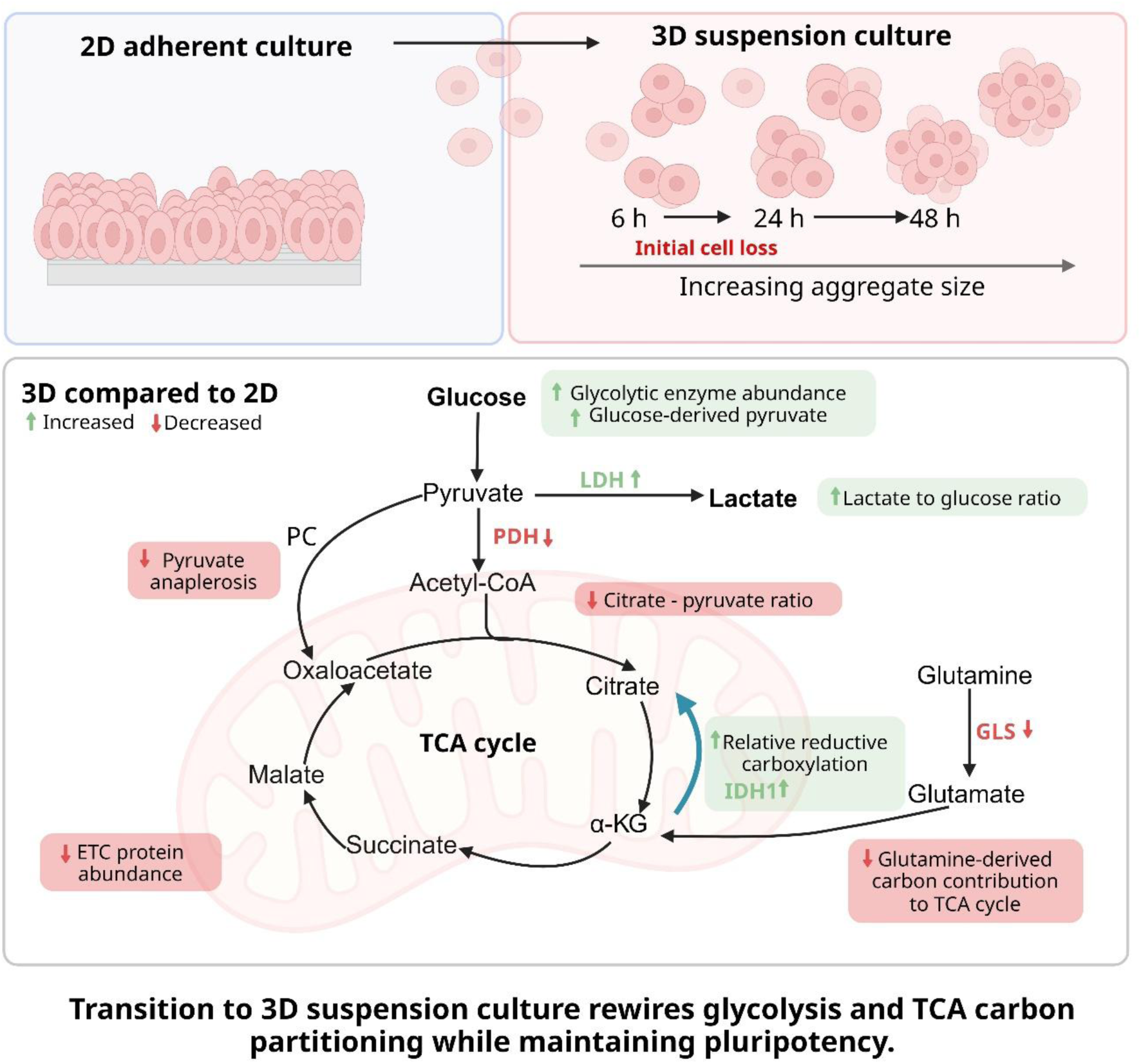

## Introduction

Degenerative disorders such as Parkinson’s disease and type 1 diabetes are characterized by the progressive loss of specific cell types, creating a critical need for cell-replacement therapies. Transplantation of human donor tissue has demonstrated the feasibility of cell replacement as a therapeutic and potentially curative approach (Lindvall et al., 1989; Shapiro et al., 2000). However, the availability of donor material is limited, highlighting the need for alternative cell sources.

Human pluripotent stem cells (hPSCs) provide a scalable alternative, enabling the generation of patient-relevant cell types without reliance on donor tissues (Takahashi et al., 2007; Thomson et al., 1998). Differentiation protocols have been optimized to generate therapeutically relevant cell types (Balboa et al., 2022; Doi et al., 2014; Kim et al., 2021; Nolbrant et al., 2017; Pagliuca et al., 2014; Rezania et al., 2014), and an increasing number of studies are advancing toward clinical translation (Chang et al., 2025; Kirkeby et al., 2025; Reichman et al., 2025; Sawamoto et al., 2025; Tabar et al., 2025). Despite this progress, billions of cells are required to make cell-replacement therapy accessible as an off-the-shelf product (Kirkeby *et al*., 2025). To address this demand, scalability and purity are key considerations in the development of processes for expanding and differentiating hPSCs (Cyrys and Zweigerdt, 2026). Moreover, because hPSC expansion is one of the most expensive steps in this process (Celiz et al., 2014; Chen et al., 2011; Kuo et al., 2020), there is a clear motivation to optimize culture conditions for efficient and cost-effective cell production.

Traditionally, hPSCs are cultured on matrix-coated tissue culture plastics (2D culture), due to their anchorage-dependent nature. While 2D culture systems are useful for small-scale differentiation and drug-screening, 3D suspension culture offers a scalable and controlled platform (Cyrys and Zweigerdt, 2026). Therefore, the transition from 2D to 3D suspension represents a key step toward clinical manufacturing. In suspension culture, hPSCs spontaneously form aggregates, whereas in 2D they are maintained as monolayers. Passage and inoculation protocols for both 2D and 3D systems typically require the addition of a Rho-associated protein kinase inhibitor (ROCKi) to improve cell survival (Watanabe et al., 2007). However, substantial cell loss is still observed during the transition from 2D to 3D culture (Kropp et al., 2016; Steiner et al., 2010). Although ROCKi supports survival during passaging, the observed cell loss suggests that the transition is associated with poorly understood molecular adaptations and requires further optimization.

Molecular changes associated with the transition from adherent monolayers to 3D suspension cultures have been reported in other cell systems (Kalender et al., 2022; Rybkowska et al., 2023), whereas few studies have examined these effects in the pluripotent state. Among these, studies comparing hPSCs cultured in 2D and 3D suspension cultures have primarily focused on molecular changes in cell-cell adhesion, particularly related to E-cadherin and WNT-signaling (Azarin et al., 2012; Konze et al., 2014).

Beyond these molecular comparisons, much of the existing work has focused on developing protocols for hPSC expansion in suspension cultures (Manstein et al., 2019; Manstein et al., 2021) and on optimizing suspension culture systems in small-scale and scalable bioreactors (Kriedemann et al., 2024a; Kropp *et al*., 2016). Although these studies primarily focus on long-term expansion in suspension culture, some include adherent 2D starting material in their analyses, demonstrating changes in cell-cell and cell-matrix adhesion molecules between the 2D and suspension cultures. Additionally, evidence points to alterations in metabolic pathways including glycolysis and nucleotide metabolism (Kropp et al., 2016; Silva et al., 2015). However, it remains unclear whether, and to what extent, the metabolic state of hPSCs changes during the transition from 2D monolayer to 3D suspension culture.

Here, we systematically investigated molecular changes associated with the transition from 2D to 3D suspension culture in hPSCs using a multi-omics approach. Proteomic pathway analysis identified alterations in glucose metabolism, prompting subsequent stable-isotope tracing metabolomic analyses. We found that 3D suspension–cultured hPSCs exhibit increased formation of glucose-derived pyruvate and rewired glucose carbon partitioning within the TCA cycle, including reduced pyruvate carboxylase-dependent anaplerosis. Glutamine contribution to the TCA cycle was reduced, whereas the relative reductive carboxylation of glutamine-derived carbon was increased. These findings demonstrate that suspension culture induces metabolic rewiring in hPSCs, providing a framework for further optimization of scalable culture systems.

## Results

### Establishment and validation of 2D and 3D hPSC culture systems for comparative analysis

To establish a dynamic 6-well plate-based 3D suspension culture system and enable comparative characterization of hPSCs in 2D and 3D culture conditions, we first adapted hPSC lines (hESC and hiPSC) to iPS-Brew, a pluripotency medium that supports culture in both formats. Karyotyping demonstrated chromosomal integrity and flow cytometry confirmed expression of pluripotency markers following adaptation (Supplementary Figure 1A–C). Furthermore, we identified optimal seeding density of 0.25×10⁶ cells/ml for 3D suspension culture (Supplementary Figure 1D–H). For all experiments, hPSCs were thawed and propagated in 2D culture for two to four passages prior to the experimental setup (Figure 1A). hPSCs from 2D and 3D cultures were sampled after 6, 24, and 48 hours for downstream analyses. Cell morphology evolved over time, from newly seeded cells (6 hours) and immature aggregates to more defined, round aggregates at 24 and 48 hours (Figure 1B). Similarly, in 2D cultures, single cells initially attached to the coated surface and began to spread (6 hours), followed by colony formation and expansion (Figure 1B). Consistent with visual observations, aggregate size increased over time (Figure 1C). An increased number of free-floating cells was observed in 3D cultures relative to 2D cultures at the 24-hour time point (Figure 1D). Consistent with this observation, the fold change in cell number between 0 and 24 hours was significantly lower in 3D suspension cultures (Figure 1E). Between 24 and 48 hours, fold changes indicated a higher proliferation rate in adherent cultures. Mean cell viability remained high across conditions (Figure 1F), although greater variability and slightly lower viability were observed in 3D suspension compared to 2D cultures. >90% of hPSCs co-expressed SOX2 and OCT3/4 in both culture systems, but the percentage of double-positive cells was slightly lower and more variable in 3D suspension cultures compared to 2D across all time points (Figure 1G).

**Figure 1.**
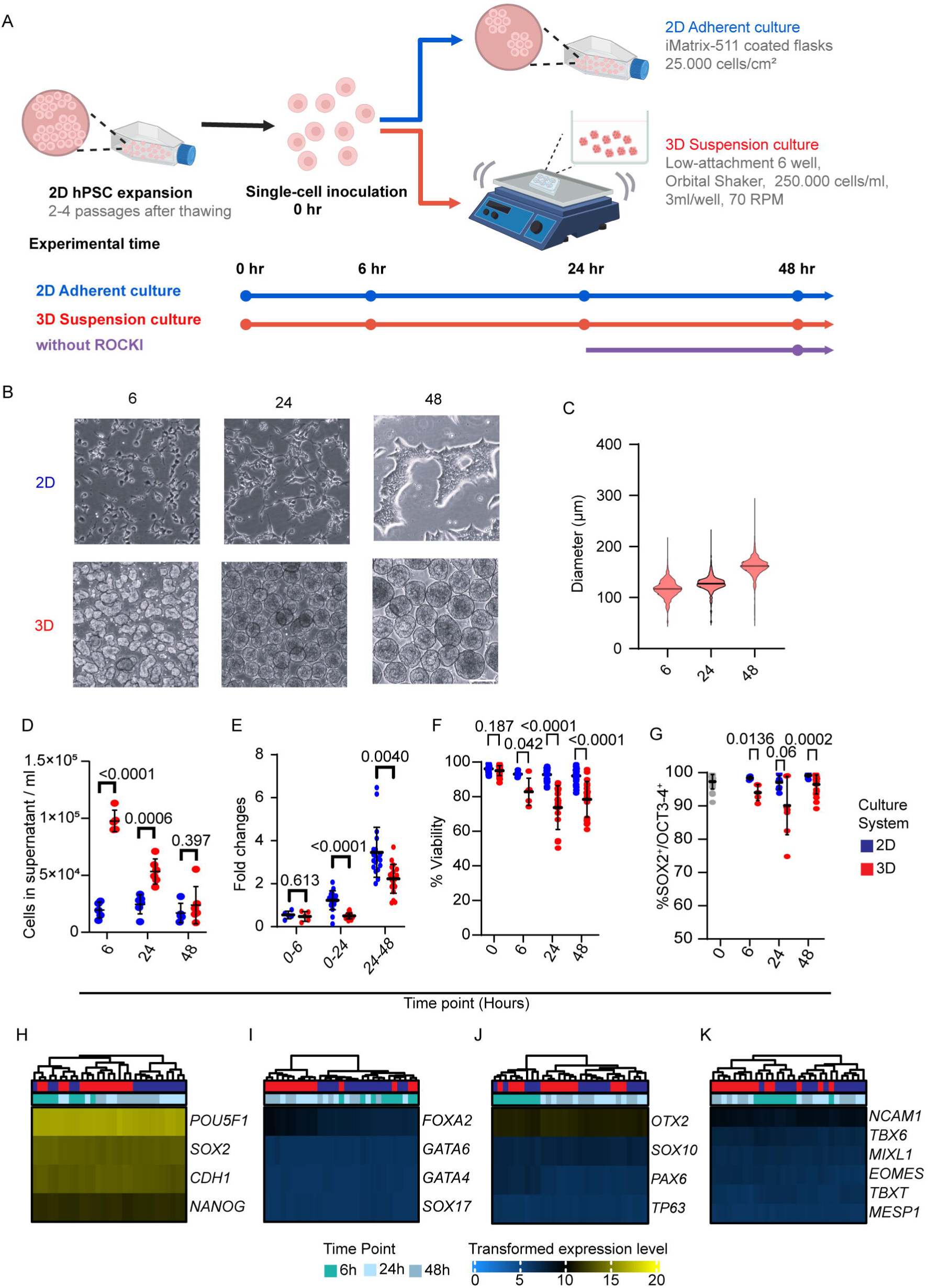
hPSCs maintain pluripotency despite differential growth in 2D and 3D culture. A) Schematic overview of the experimental design. Following expansion in 2D adherent culture, hPSCs were dissociated into single cells and inoculated into either 2D adherent or 3D suspension culture. Samples were collected at 6, 24, and 48 hours post-inoculation. The effect of ROCK inhibitor (ROCKi) withdrawal in 3D suspension culture after 24 hours was assessed in selected experiments, as shown in Supplementary Figure 5.B) Representative bright-field images of 2D and 3D cultures at each time point. Scale bar: 200 μm. C) Aggregate diameter (μm) in 3D suspension culture at each time point, measured using the BIOREP islet counter. D–G) Characterization of hPSCs over time in 2D and 3D suspension culture. D) Cell count in the supernatant collected during medium change showed as cells/ml. E) Cell growth expressed as fold changes at 0–6, 0–24, and 24–48 hours. F) Cell viability (%) at inoculation (0 hours) and the three time points. G) Pluripotency assessed by the percentage of cells co-expressing SOX2 and OCT3/4, measured by flow cytometry at inoculation and at each time point. Data is presented as mean ± SD with individual biological replicates shown. N = 5–22 depending on the measurement. Statistical comparisons were performed using Welch’s t-test; p-values are shown above the comparison. H–K) Bulk RNA sequencing analysis of lineage-specific gene expression, shown as heatmaps of transformed expression levels, categorized by culture system (2D vs. 3D) and time point (6, 24, and 48 hours). H) Pluripotency, I) Endoderm, J) Ectoderm, and K) Mesoderm associated genes. N = 5 biological replicates. For validation of the suspension culture and cell line see supplementary Figure 1.

To explore the molecular changes occurring in hPSCs cultured under 2D and 3D suspension conditions, we performed bulk RNA-sequencing and proteomics at 6, 24, and 48 hours post-inoculation. Given the small but significant differences in SOX2 and OCT3/4 expression, RNA-sequencing was used to assess potential spontaneous differentiation. Expression of pluripotency-associated genes (*POU5F1/OCT3/4, SOX2, CDH1, NANOG*) and lineage markers for endoderm (*FOXA2, GATA6, GATA4, SOX17*), ectoderm (*OTX2, SOX10, PAX6, TP63*), and mesoderm (*NCAM1, TBX6, MIXL1, EOMES, TBXT, MESP1*) was evaluated (Figure 1H–K). This analysis confirmed maintenance of pluripotency with no evidence of differentiation across both culture systems at all time points. Furthermore, to assess whether the transition from 2D to 3D suspension culture was associated with changes in the pluripotent state, we examined the expression of genes associated with the naïve pluripotent state (*VENTX, TFAP2C, KLF4, DPPA3, FGF4, DPPA5, TBX3, KLF17*) (Supplementary Figure 1I-J). The expression patterns of these genes provided no indication of a shift towards a naïve pluripotent state. Overall, these data show that while 2D and 3D cultures differ in growth and morphology, primed pluripotency was maintained with no evidence of spontaneous differentiation, supporting the use of these systems as a robust foundation for comparative molecular analyses of hPSCs cultured in 2D and 3D systems.

### Culture format drives molecular variation in hPSCs

Using the transcriptomic and proteomic datasets, we examined the major sources of variation between samples. Principal component analysis (PCA) of both datasets demonstrated clear separation of samples along PC1 according to culture format, accounting for 34.9% and 54% of the variance in the proteomic and transcriptomic datasets, respectively (Figure 2A, B). In contrast, sampling time point drove variation along PC2, explaining 14.1% and 27% of the variance in the proteomic and transcriptomic datasets, respectively. These results indicate that culture format is the dominant driver of molecular variation, while time point contributes to a smaller extent as cultures progress over time. We observed a high degree of overlap between genes identified in the proteomic and transcriptomic datasets (5,212 out of 5,621 proteins) (Figure 2C). However, only 32 shared differentially expressed genes were identified across both datasets (Figure 2D). The abundances of these shared hits showed moderate correlation (Figure 2D, Supplementary Figure 2A, B), consistent with previous reports, suggesting post-transcriptional modifications and turnover rates may contribute to the discrepancy (Konze *et al*., 2014; Koussounadis et al., 2015; Nevedomskaya et al., 2022; Vogel and Marcotte, 2012).

**Figure 2.**
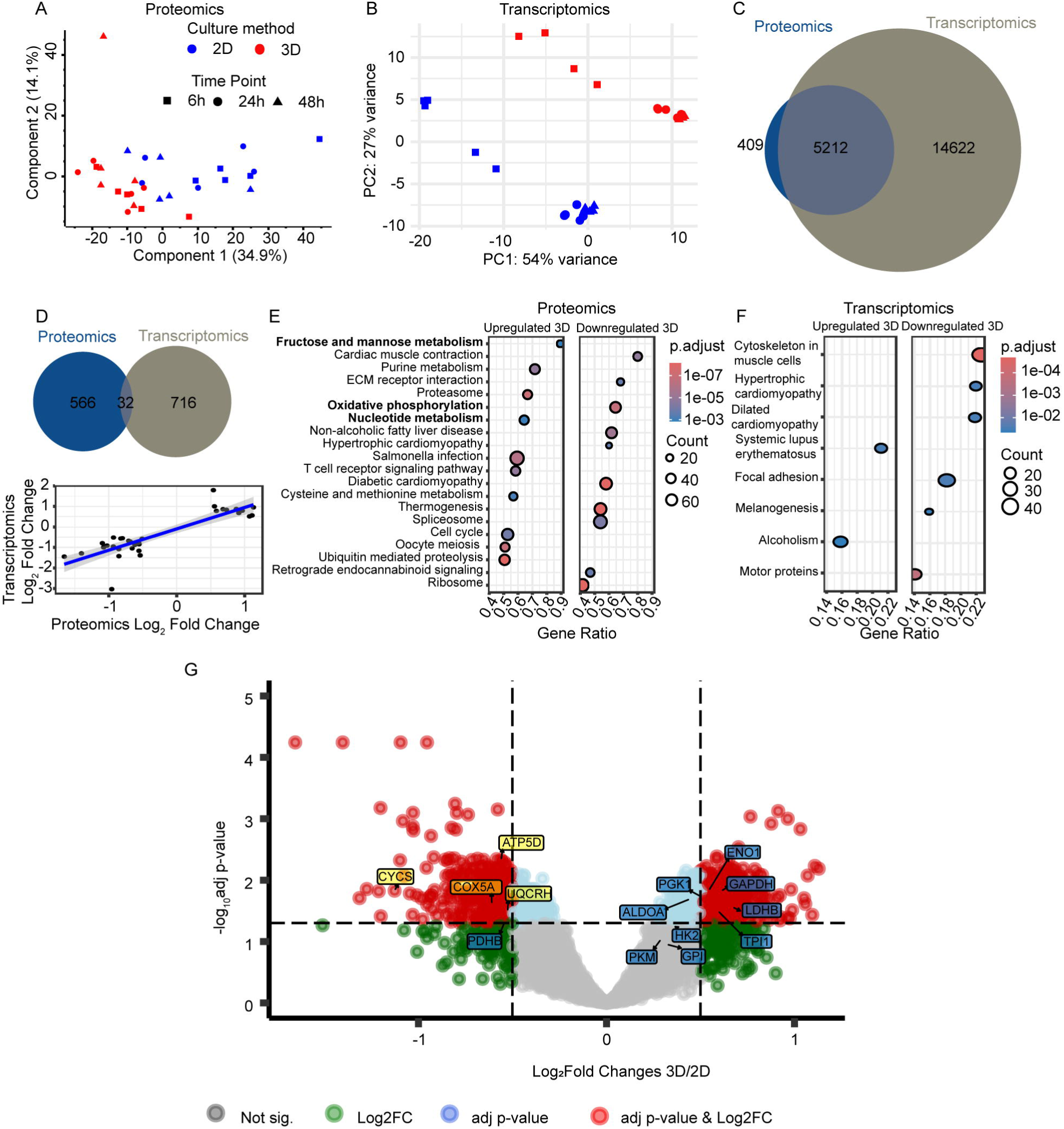
Molecular changes between 2D and 3D suspension cultured hPSCs. A) Principal component analysis (PCA) of proteomic data, showing PC1 and PC2. B) PCA of transcriptomic data, showing PC1 and PC2. In both A) and B), data points are colored by culture format and shaped by time point. C) Venn diagram of the overlap between total detected proteins and transcripts. D) Venn diagram shows the overlap between differentially expressed proteins and genes at 48 hours (top), and correlation of Log_2_ fold changes (FC) between proteomics and transcriptomics for the overlapping hits at 48 hours (bottom). The blue line represents a linear regression fit. E-F) KEGG pathway enrichment analysis at 48 hours. Dot size reflects gene ratio, and color indicates adjusted p-value. E) Proteomic and F) Transcriptomic. G) Volcano plot of proteomic data at 48 hours comparing 3D suspension versus 2D culture. Glycolytic enzymes are highlighted in blue and oxidative phosphorylation (OxPhos) subunits in yellow, including Complex I (NDUFA2), Complex III (UQCRH), Complex IV (COX5A), Cytochrome C (CYCS), and ATP synthase (ATP5D). Dashed vertical lines indicate a Log_2_ fold change cutoff > 0.5 (FC >|1.41|). The dashed horizontal line indicates an adjusted p-value threshold of 0.05 on the -log_10_ scale. For proteomics FDR = 0.05. Data represents N = 5 biological replicates. For pathway analysis at 6 and 24, see supplementary Figure 2.

### Proteomic analysis revealed culture format–dependent changes in glycolytic enzyme abundance

Next, we performed gene set enrichment analysis (GSEA) using KEGG pathways to identify the biological pathways associated with the molecular variation across culture formats. KEGG pathway analysis, performed independently on the transcriptomic and proteomic datasets, revealed shared changes related to cytoskeletal reorganization and adhesion pathways (e.g. TUBA1A, BCAM, SDC2 and VIM, in both; CLDN6 and TUBB in proteomic data; and *MYL9* in transcriptomic data) (Figure 2E, F). Similar enrichment of cytoskeletal and adhesion pathways was observed at 6 and 24 hours (Supplementary Figure 2). While cytoskeletal reorganization and adhesion pathways showed consistent changes across both datasets, the proteomic data additionally revealed enrichment of pathways related to glucose metabolism and mitochondrial oxidative phosphorylation (Figure 2E; full list is found in supplementary data). Specifically, proteomic analysis showed significantly increased abundance of glycolytic enzymes in 3D suspension cultures (Figure 2G), suggesting a shift in glucose metabolism associated with the transition from 2D to 3D culture conditions. In contrast, proteins associated with the electron transport chain (ETC) were downregulated in 3D suspension cultures compared to 2D cultured hPSCs (Figure 2E). The proteomic data further indicated opposing changes in abundance of lactate dehydrogenase (LDH) and pyruvate dehydrogenase (PDH), which convert pyruvate to lactate and acetyl-CoA, respectively (Figure 3A). Western blot analysis confirmed increased LDH and decreased PDH protein abundance in 3D suspension cultures (Figure 3B-D). Together, these findings show that the abundance of key metabolic enzymes is markedly altered upon the transition from adherent to suspension culture conditions suggesting a profound metabolic rewiring induced by the culture format.

**Figure 3.**
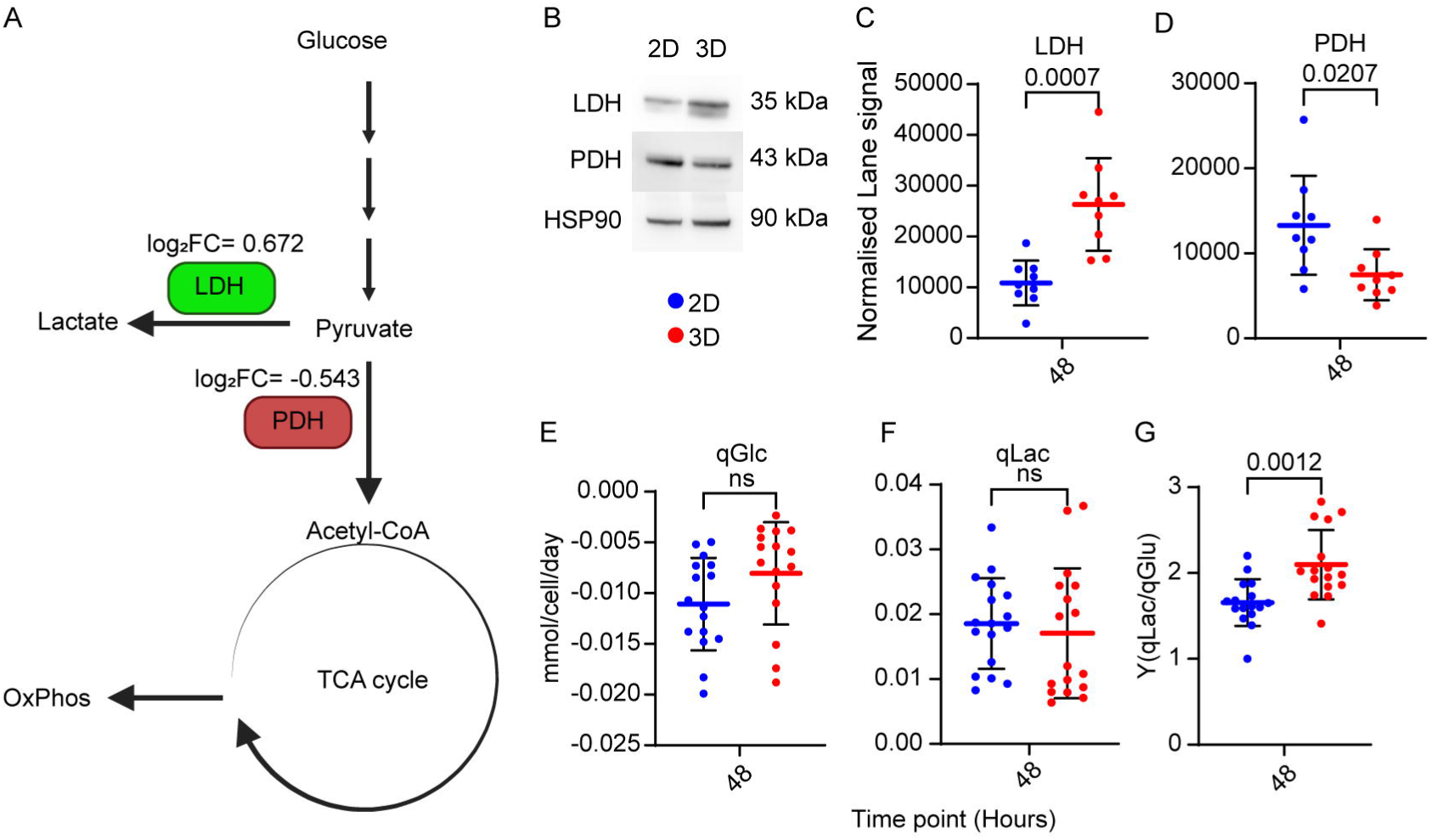
Differential abundance of LDH and PDH and elevated lactate to glucose ratio support increased glycolysis in 3D cultured hPSCs. A) Schematic illustration of glycolysis, the TCA cycle, and oxidative phosphorylation (OxPhos), highlighting the regulation of LDH (green, upregulated) and PDH (red, downregulated) in 3D suspension culture, with log_2_FC from proteomics. B) Representative western blots of LDH, PDH, and HSP90 (loading control). C) LDH and D) PDH abundance quantification from western blotting. N = 9. E-G) spent-medium analysis. E) Consumed glucose (qGlc) and F) lactate production (qLac), G) Lactate yield from glucose (Y(qLac/qGlc)). N = 17. Data presented as mean ± SD with individual biological replicates. Statistical analysis was performed using Welch’s t-test, p-values are shown above. For the 24-hour time point see supplementary Figure 3, and for assessment of STB, cell line and medium see supplementary Figure 4.

### Spent growth media analysis confirms culture format–dependent metabolic differences in hPSCs

Following the observed upregulation of glycolytic enzymes, we performed spent media analyses to quantify glucose consumption and lactate production as a functional measure of glycolytic flux toward lactate. Glucose consumption (qGlc) was significantly higher in 3D compared to 2D after 24 hours (Supplementary Figure 3A), but not at 48 hours (Figure 3E). Lactate production (qLac) did not differ significantly between 2D and 3D suspension cultures (Figure 3F and Supplementary Figure 3A). However, by evaluating the yield-specific ratio qLac/qGlc (YqLac/qGlc) (Figure 3G and Supplementary Figure 3A), an indicator of the relative glucose metabolism to lactate, we found significantly increased YqLac/qGlc in 3D suspension cultures, a result similar to previous observations in stirred-tank bioreactor systems (Silva *et al*., 2015). This finding supports the notion that hPSCs in 3D suspension exhibit increased reliance on glycolytic metabolism, consistent with the proteomic data (Figure 2G).

To determine whether these findings were generalizable beyond the specific conditions employed, we assessed YqLac/qGlc as well as LDH and PDH abundance in a human induced pluripotent stem cell (iPSC) line. We found consistent trends, with increased YqLac/qGlc in 3D suspension cultures and corresponding LDH and PDH protein expression patterns confirmed by western blotting (Supplementary Figure 4A–D). Consistent results were also observed when hPSCs were inoculated into a stirred-tank bioreactor (DASbox) (Supplementary Figure 4E–H). Furthermore, similar patterns in YqLac/qGlc and LDH and PDH expression were observed when using a different pluripotency medium (NutriStem® hPSC XF Medium) (Supplementary Figure 4I-L). These findings demonstrate that the observed metabolic shift is associated with the transition from 2D to 3D suspension cultures, independently of cell line, medium, and suspension format, and is reflected in both the protein signature and functional metabolic output.

## 3D suspension culture enhances glucose-derived pyruvate formation and rewires TCA cycle glucose carbon partitioning

In the tricarboxylic acid (TCA) cycle carbon is oxidized, generating reducing agents (NADH and FADH_2_) for oxidative phosphorylation (OxPhos), but TCA intermediates also serve as precursors for biosynthesis, as well as for chromatin remodelling and post-translational modifications (Vander Heiden et al., 2009). We identified minimal changes in the abundance of TCA enzymes, with downregulation of malate dehydrogenase 2 (MDH2) in 3D suspension being the exception. However, we observed differential abundance of glutaminase (GLS), isocitrate dehydrogenase 1 (IDH1), and malate dehydrogenase 1 (MDH1) (Figure 4A), related to cytosolic processing of glutamine and TCA metabolites. To further investigate the differences between 2D and 3D cultures with respect to cellular metabolism, we conducted stable isotope tracing metabolomic analyses using uniformly labelled glucose ([U-^13^C]-Glucose) and glutamine ([U-^13^C]-Glutamine), focusing on their contribution to TCA cycle metabolites. For this study, we decided to include a 3D condition without ROCKi from 24 to 48 hours, as ROCKi has been reported to modulate metabolism (Matsumoto et al., 2022; Vernardis et al., 2017). However, by assessing Y(qLac/qGlc) between 3D cultured hPSCs with or without ROCKi, we did not observe a significant influence of ROCKi supplementation (Supplementary Figure 5A–I). The tracer metabolomic analysis revealed substantial differences between 2D and 3D suspension-cultured hPSCs, and minor differences between 3D suspension-cultured hPSCs with or without ROCKi. However, ROCKi slightly increased the magnitude of differences between 2D and 3D suspension cultures when added beyond 24 hours (Supplementary Figure 5J–N).

**Figure 4.**
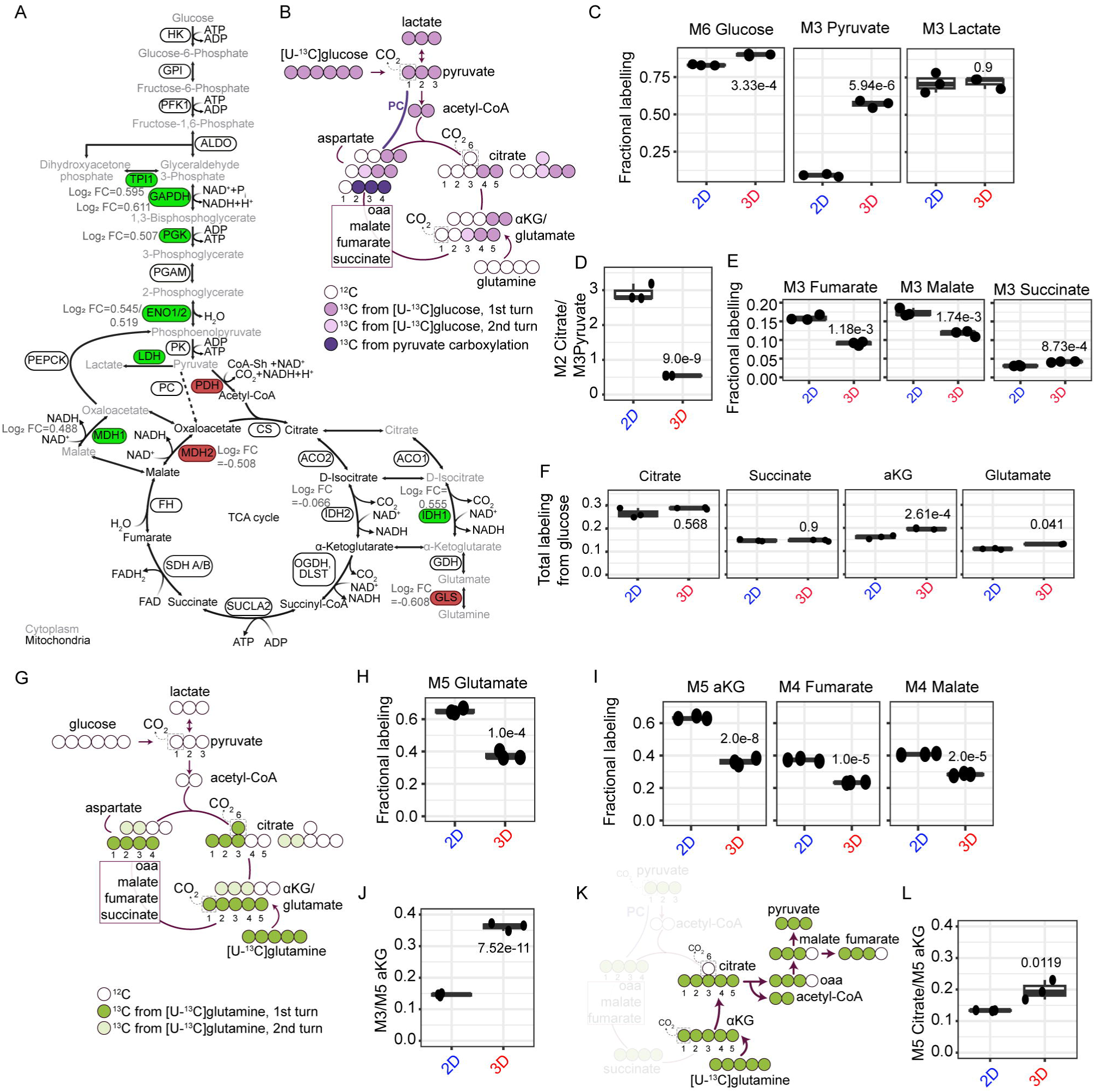
Stable-isotope tracer metabolomic analysis revealed altered glucose and glutamine carbon partitioning in 3D suspension-cultured hPSCs. **A)** Schematic overview of glycolysis and the tricarboxylic acid (TCA) cycle displaying metabolites and enzymes, differentially expressed enzymes are highlighted in green and red, indicating significantly increased and reduced protein abundance, respectively, in 3D suspension cultures with corresponding log₂ fold change (Log₂ FC) values. HK (Hexokinase), GPI (Glucose-6-phosphate isomerase), PFK (phosphofructokinase), ALDO (aldolase), TPI1 (triose phosphate isomerase), GAPDH (glyceraldehyde 3-phosphate dehydrogenase), PGK (Phosphoglycerate kinase), PGAM (phosphoglycerate mutase), ENO (enolase), PK (Pyruvate kinase), LDH (lactate dehydrogenase), PDH (pyruvate dehydrogenase), CS (Citrate synthase), ACOS1/2 (aconitase), IDH (isocitrate dehydrogenase), OGDH/DLST (α-ketoglutarate dehydrogenase complex), SUCLA (Succinyl-CoA synthetase), SDH (succinate dehydrogenase), FH (fumarase), MDH (malate dehydrogenase), GLS (glutaminase), and GDH (glutamate dehydrogenase). B) Schematic representation of carbon transition from [U-^13^C]-glucose into pyruvate, lactate and TCA metabolites. Fully labelled glucose (M6) generates M3 pyruvate through glycolysis, which can enter the TCA cycle via PDH or pyruvate carboxylase (PC), generating distinct isotopologue distributions in downstream metabolites. C–F) Isotopic labeling of metabolites following [U-^13^C]-glucose labelling in 2D and 3D cultures. C) Fractional labelling of M6 glucose, M3 pyruvate, and M3 lactate. D) Ratio of M2 citrate to M3 pyruvate labelling as an indicator of glucose-derived carbon entry into the TCA cycle. E) Fractional labelling of second round TCA intermediates M3 fumarate, M3 malate, and M3 succinate. F) Total ^13^C labelling from glucose in citrate, succinate, α-ketoglutarate (αKG), and glutamate. G) Schematic representation of carbon transition from [U-^13^C]-glutamine. Glutamine replenishes the TCA cycle through αKG. The first round generates M5 αKG and M4 isotopologues of downstream intermediates through oxidative TCA metabolism. H–J) Isotopic labeling of metabolites following [U-^13^C]-glutamine labelling in 2D and 3D cultures. H) Fractional labelling of M5 glutamate. I) Fractional labelling of first round M5 αKG, M4 fumarate, and M4 malate. J) Ratio of M3 αKG to M5 αKG. K) Schematic representation of reductive carboxylation from [U-^13^C]-glutamine-derived αKG and M5 citrate. L) Ratio of M5 citrate to M5 αKG, used as a measure of reductive TCA metabolism from glutamine. n = 3 technical replicates; data presented as mean with individual replicates. Statistical significance was assessed using one-way ANOVA and Tukey’s HSD test, p-values are shown for each comparison. See also supplementary Figure 3 for 24 hours, Supplementary Figure 5, for ROCKi assessment and Supplementary Figure 6.

[U-^13^C]-Glucose is converted through glycolysis to pyruvate and lactate, both labelled at all three carbons (M3 isotopologues) (Figure 4B). At 48 hours after inoculation, we observed a significantly higher fractional enrichment in intracellular M6 glucose and M3 pyruvate in 3D suspension compared to adherent cultures (Figure 4C). The increased M6 glucose enrichment suggested a potentially higher glucose uptake, whereas the elevated M3 pyruvate/M6 glucose ratio indicated an increase in relative glucose metabolism towards pyruvate in 3D suspension compared to 2D culture (Supplementary Figure 6). Despite the higher labelling in pyruvate in 3D suspension cultures, we observed comparable incorporation of ^13^C into lactate between the two culture conditions (Figure 4C), which was also reflected by the qLac measured in the spent culture media (Figure 3F). Consistent with the 48-hour findings, a similar pattern of metabolite labelling was observed at the 24-hour time point (Supplementary Figure 3D, E).

Pyruvate continues to the TCA cycle via PDH-mediated conversion to acetyl-CoA, generating TCA metabolites labeled in two carbon atoms by the end of the first round of the cycle (M2 isotopologues) (Figure 4B). M2 labeling of TCA cycle metabolites was significantly increased in 3D suspension compared to 2D cultured hPSCs at both 24 and 48 hours (Supplementary Figure 6 and 7). However, the M2 citrate/M3 pyruvate ratio was reduced, indicating that the relative pyruvate entry through the PDH route was decreased in 3D suspension cultures (Figure 4D), which is supported by proteomic and western blot data revealing downregulation of PDH in 3D suspension.

To better understand the impact of glucose as a carbon source for TCA cycle metabolites, we calculated the total carbon contribution of glucose to the intermediates. At 24 hours, glucose contribution to citrate, α-ketoglutarate (α-KG), glutamate, and succinate was increased in 3D cultures, whereas its contribution to fumarate and malate was reduced (Supplementary Figure 3F, G). By examining the labelling patterns at both 24 and 48 hours, we observed that the M3 fumarate and M3 malate (Figure 4E and Supplementary Figure 3H) were instead enriched in adherent hPSCs. Although these isotopologues can arise during the second round of the TCA cycle (Figure 4B), their selective enrichment in the absence of a corresponding increase in M3 succinate (Supplementary Figure 3H, and Supplementary Figure 6) is consistent with greater pyruvate carboxylase-mediated anaplerosis in adherent cells (Figure 4B).

By 48 hours, glucose contribution to citrate and succinate had equalised between the two culture conditions, whereas α-KG and glutamate remained more highly labelled in the 3D suspension cultures (Figure 4F). The selective enrichment of the α-KG/glutamate node, together with the reduced labelling of the distal C4 pools suggests potentially altered carbon retention and exchange, as well as reduced dilution by unlabelled carbon substrates around this metabolic node. Collectively, these data indicate that 3D suspension culture increases the relative glucose metabolism to pyruvate and redistributes glucose carbon incorporation across the TCA cycle. In addition to glucose and pyruvate, glutamine is an important carbon source feeding into the TCA cycle (Kafkia et al., 2026; Tohyama et al., 2016).

### 3D suspension culture limits glutamine carbon entry into the TCA cycle while promoting reductive carboxylation

Glutamine is converted to glutamate by GLS (Tohyama *et al*., 2016), which we found to be significantly downregulated in 3D suspension–cultured hPSCs (Figure 4A). [U-^13^C]-glutamine transfers five ^13^C atoms to glutamate (M5 isotopologues) (Figure 4G), and our metabolomics data showed a significant enrichment of M5 glutamate in 2D compared to 3D suspension cultures (Figure 4H), supporting our proteomic findings. M5 glutamate can be converted to M5 α-KG, entering the first round of the TCA cycle and transferring four ^13^C atoms to downstream TCA metabolites (M4 isotopologues) (Figure 4G). Consistently, we observed significantly higher fractional labelling of M4 TCA metabolites in adherent compared to 3D suspension cultured hPSCs (Figure 4I).

In the second round of the TCA cycle, M4 citrate transfers one carbon to CO₂, resulting in M3 α-KG and M3 glutamate, as well as M2 isotopologues of succinate, fumarate and malate (Figure 4G). The second-round tracing of [U-^13^C]-glutamine showed the opposite pattern to the first round with significantly increased incorporation of ^13^C atoms in 3D suspension cultures compared to adherent cultures (Supplementary Figure 6). Moreover, the ratio of M3 α-KG to M5 α-KG was likewise significantly elevated in 3D suspension cultured hPSCs (Figure 4J). This pattern may reflect increased TCA cycling, reduced dilution by alternative sources, and/or reduced cataplerotic withdrawal as additional characteristics of the 3D suspension cultured hPSCs. In line with this, total glutamine contribution to the TCA cycle metabolites was higher in the adherent cultures (Supplementary Figure 6).

Notably, we also observed greater redirection of glutamine-derived carbon towards reductive carboxylation in the 3D suspension cultures, as indicated by elevated M5 citrate and the M5 citrate/ M5 α-KG ratio (Figure 4K, L). This shift is likely linked to the increased protein abundance of IDH1 in 3D suspension cultures (Figure 4A). IDH1 is the cytosolic analogue of IDH2, which was not differentially expressed at the 48-hour time point. The reductive carboxylation of glutamine-derived α-KG can support citrate and acetyl-CoA production for lipid biosynthesis and histone acetylation (Kafkia et al., 2026; Metallo et al., 2012). Together, our findings reveal a broad metabolic reorganization in the 3D suspension cultures, characterized by increased glycolytic conversion of glucose to pyruvate and altered TCA cycle carbon partitioning, including reduced pyruvate carboxylase-dependent anaplerosis and glutamine contribution, and increased reductive utilization of glutamine-derived carbon.

## Discussion

The clinical translation of cell replacement therapies requires scalable and tightly controlled culture systems. To meet this need, hPSC culture has increasingly transitioned toward 3D suspension culture systems, such as stirred-tank bioreactors and Vertical-Wheel bioreactors, which offer improved scalability compared to traditional adherent systems (Cohen et al., 2023; Cuesta-Gomez et al., 2023; Kriedemann et al., 2024b; Manstein *et al*., 2021). Although 3D suspension systems are increasingly adopted for scalable stem cell culture, traditional adherent (2D) formats remain widely used (Cyrys and Zweigerdt, 2026). Defining how these culture conditions differentially shape hPSC metabolism and function is critical for achieving robust control of expansion and differentiation. To this end, we established a small-scale culture system using 6-well low-attachment plates, similar to previously described systems (Konze *et al*., 2014), and validated its performance prior to conducting an unbiased multi-omics analysis of adherent (2D) and 3D suspension–cultured hPSCs. We observed a substantial reduction in the number of cells in suspension cultures after 24 hours in culture, as reflected by the observation of a larger number of free-floating cells at this time point. This observation is consistent with previous reports (Borys et al., 2021; Kropp *et al*., 2016; Manstein *et al*., 2021; Steiner *et al*., 2010; Ullmann et al., 2024). We interpret this initial cell loss as potentially reflecting an adaptation phase following transfer to the suspension cultures (Ullmann *et al*., 2024). Despite this reduction in cell numbers, pluripotency was conserved.

Our multi-omics analysis demonstrated that the culture system (2D or 3D) was the primary driver of the observed biological changes as opposed to timing since inoculation, underscoring the relevance of studying the transition between these culture formats. Pathway analysis revealed downregulation of “Cell adhesion molecules” and “ECM–receptor interaction” in suspension cultures, consistent with previous reports (Konze et al. 2014; Azarin et al. 2012; Ullmann et al. 2024). However, when we examined the overlap between regulated proteins reported in (Konze *et al*., 2014) and our data, only a partial overlap was observed (e.g. CNN1, PODXL, PKP2, TPM1, LMNA). This could be a consequence of differences in hPSC lines, culture conditions, and time points between studies.

In addition to cytoskeletal changes, our proteomic GSEA revealed enrichment of glucose metabolism pathways and downregulation of oxidative phosphorylation in 3D suspension cultures, reminiscent of the Warburg effect described in cancer cells (Vander Heiden *et al*., 2009). Cancer cells preferentially convert glucose to lactate despite sufficient oxygen availability to support rapid proliferation. In the present study, hPSCs were cultured in CO_2_-controlled incubators, thus oxygen levels in the incubator were assumed to be close to atmospheric levels. Cellular metabolism is highly dynamic and adapts to cellular demands and differentiation state (Jackson and Finley, 2024). Similarly, dynamic metabolic regulation has been described across pluripotent stem cell stages, transitioning from a more bivalent metabolic profile in naïve hPSCs to a predominantly glycolytic metabolism in primed hPSCs, and subsequently rewiring upon differentiation to match lineage-specific metabolic requirements (Kafkia *et al*., 2026; Van Nerum et al., 2025). α-KG is found to regulate the pluripotent stages through DNA and histone demethylations. In naïve mouse ESCs, α-KG is found to maintain pluripotency, whereas α-KG-dependent demethylation promotes differentiation in primed hPSCs (Carey et al., 2015; TeSlaa et al., 2016). Furthermore, the transition to 3D suspension culture has been reported to support the maintenance of naïve pluripotency (Rohani et al., 2020). In our system, glucose-derived carbons contributed to increased α-KG labeling in suspension cultures. We did not observe changes in the expression of naïve pluripotency markers between culture systems or time points, suggesting that the culture conditions alone are not driving the cells towards a more naïve state.

YqLac/qGlc is widely used as a non-invasive in-process parameter reflecting metabolic activity and culture performance but is rarely directly compared between adherent and suspension cultures. We observed a significantly increased YqLac/qGlc in suspension cultures compared to adherent cultures, a trend consistent across cell lines, culture systems, and various hPSC media conditions. Silva and coworkers similarly reported increased YqLac/qGlc and reduced growth in bioreactor cultures compared to adherent conditions, attributing lactate production partly to non-glucose carbon sources when YqLac/qGlc is >2 (Silva *et al*., 2015). In the current study, YqLac/qGlc is much closer to 2, the theoretical ratio if only glucose contributes to lactate production. Increased YqLac/qGlc ratio and an elevated M3 pyruvate/M6 glucose ratio support the interpretation that 3D suspension cultures are more glycolytic. In contrast, Kropp and coworkers reported the opposite trend, potentially reflecting differences in time point or feeding strategy (Kropp *et al*., 2016). Consistent with our results, other RNA-sequencing-based studies comparing 2D and 3D suspension cultured hPSCs, (Konze *et al*., 2014; Kropp *et al*., 2016; Ullmann *et al*., 2024) did not identify regulation of metabolic pathways, suggesting that these changes may not be driven at the transcriptional level, but may instead result from downstream regulation and/or post-transcriptional modifications. Konze and coworkers also performed proteomics, but did not report enrichment of glucose or TCA pathways (Konze *et al*., 2014). However, GAPDH, a key glycolytic enzyme commonly used as a housekeeping protein, was differentially regulated in their dataset, indicating that metabolic changes may have been present but not captured at the pathway level. Together, these observations point to a functional metabolic shift during the transition to suspension culture, consistent with the concept of metabolic plasticity in stem cells (Folmes et al., 2012), suggesting that the culture system represents an underappreciated driver of metabolic adaptation in hPSCs.

To characterize this in greater detail, we applied stable isotope tracing metabolomics to investigate glycolytic endpoints (pyruvate and lactate) as well as TCA metabolites. [U-^13^C]-Glucose labelling confirmed increased glycolytic activity towards pyruvate in suspension culture. Glycolysis is known to maintain both pluripotency and cell proliferation (Kim et al., 2015). We further observed elevated labeling of the second round isotopologues from [U-^13^C]-Glutamine, alongside increased reductive carboxylation, linked to the elevated abundance of cytosolic IDH1. Notably, glutamine contributed more extensively to TCA metabolites in 2D cultures, whereas in suspension cultures glucose-derived carbon appeared to dominate early TCA intermediates, with glutamine contributing more prominently to downstream metabolites (α-KG, succinate, fumarate, and malate). This distribution of contributions to early or late TCA metabolites further highlights the role of glycolytic intermediates in supporting biomass production during proliferation, consistent with the concept of metabolic flexibility in hPSCs (Tanosaki et al., 2020; Varum et al., 2011).

Glutamine has been shown to be essential both for maintaining pluripotency and supporting proliferation (Marsboom et al., 2016; Tohyama *et al*., 2016). It also contributes to biomass production through cataplerosis, whereby TCA intermediates, including oxaloacetate and citrate, are exported from the mitochondria to the cytosol and converted to pyruvate or acetyl-CoA for fatty acid synthesis, for example through reductive carboxylation (Kafkia *et al*., 2026; Metallo *et al*., 2012; Tohyama *et al*., 2016). The increase in cell number observed in 2D cultures suggests an increased demand for biomass production that may be supported through cataplerosis. At the same time, we observed increased reductive carboxylation in 3D suspension cultures, a process linked to fatty acid synthesis and histone acetylation via IDH1 (Kafkia *et al*., 2026).

Together, these findings point to distinct glutamine utilization strategies between culture systems and raise the possibility that culture-dependent metabolic rewiring may influence epigenetic regulation in hPSCs (Rohani *et al*., 2020). Extending beyond hPSCs, similar metabolic adaptations have been reported in mesenchymal stem/stromal cells and cancer cells (Fabiano et al., 2026; Rybkowska *et al*., 2023; Shrestha et al., 2026). Fabiano et al. further linked cell shape, adhesion and energy metabolism (Fabiano *et al*., 2026), which has also been shown by others (Bays et al., 2017; Sousa et al., 2019).

Our study demonstrates a rapid culture-dependent metabolic rewiring following transition from adherent to suspension culture, adding to the growing evidence that hPSC metabolism is highly plastic and rapidly adapts to environmental changes. We found this pattern to be consistent across cell lines, culture systems and culture media. While the mechanisms driving this adaptation remain unclear, our pathway analysis suggests several possible drivers.

The KEGG term “HIF signaling” was enriched in 3D suspension cultures after 48 hours, largely driven by increased abundance of glycolytic proteins. HIF-1 regulates the transcription of several glycolytic enzymes (PFK, PGK, ALDO, ENO, and LDH), of which many showed increased abundance in 3D suspension cultures. In addition, HIF-1 can induce PDK-mediated phosphorylation of PDH, thereby inhibiting the conversion of pyruvate to acetyl-CoA (Kierans and Taylor, 2021; Semenza et al., 1994; Zimmer et al., 2016). Although oxygen gradients within aggregates or across the medium column may have contributed to this metabolic phenotype (Winkle et al., 2012), the relatively small aggregate size suggests minimal oxygen diffusion limitations within the aggregates. As oxygen availability and HIF-1α activity were not measured, involvement of HIF signaling remains speculative.

Our study and previous studies suggest that molecular changes in cell adhesion and cytoskeletal organization could contribute to the observed metabolic rewiring. In 3D suspension cultures we observed downregulation of the KEGG pathways “ECM-receptor interaction” and “Cell adhesion molecules”. Changes in the former may be linked to the use of an extracellular matrix coating in 2D cultures (Miyazaki et al., 2008; Rodin et al., 2010). The adhesion and cytoskeletal changes could be a possible driver for metabolic rewiring. Konze and coworkers studied the transition from 2D to 3D suspension culture and identified downregulation of proteins interacting with E-cadherin including β-catenin together with increased activity of the calpain. Cleavage of E-cadherin by calpain was found essential for the aggregate formation (Konze *et al*., 2014). In our hands, calpain was significantly increased in 3D suspension cultures in our proteomic data, with no changes in β-catenin abundance. This may influence downstream signaling pathways through regulation of protein activity rather than protein abundance alone.

E-cadherin has been shown to regulate several pathways including signaling through β-catenin and YAP/TAZ (Azarin *et al*., 2012; Kim et al., 2011; Konze *et al*., 2014). YAP and TAZ have been suggested to both be regulated and act as regulators of cell metabolism (Enzo et al., 2015; Kang et al., 2024; Song et al., 2023). Neither β-catenin nor YAP showed a significant change in our data. Therefore, future studies are warranted to assess whether these factors are involved in driving the observed metabolic rewiring.

Collectively, these observations suggest that the transition from adherent to suspension culture involves remodeling of cell adhesion and cytoskeletal organization, potentially contributing to the metabolic rewiring observed during the first 48 hours of culture. Such changes may contribute to both the metabolic rewiring and the extensive cell loss observed during the first 24 hours of the suspension culture. A better understanding of the metabolic requirements during this transition could therefore support the development of improved 3D suspension cultures. Future studies should investigate whether targeted optimization of nutrient composition or other culture parameters can improve the establishment of robust suspension cultures.

## Supporting information

Proteomic Pathway analysis, 6,24,48 hours

Transcriptomics pathway analysis, 6,24,48 hours

Proteomic differential expression analysis, 6,24,48 hours

Transcriptomics differential expression analysis, 6,24,48 hours

## Resource availability

### Lead contact

Requests for further information and resources should be directed to and will be fulfilled by the lead contact, Christian Honoré.

### Material availability

This study did not generate new unique reagents.

### Data and code availability

Proteomic and RNA-seq data have been deposited at ProteomeXchange consortium via the PRIDE partner repository with the data set identifier PXD084820 and are publicly available as of the date of publication. Raw RNA-seq data cannot be provided due to privacy, ethical, and data-protection restrictions related to the cell line and is instead provided as a DESeq2 object. This paper does not report original code.

## Acknowledgements

We thank Martin Hey-Mogensen and Jens Frey Halling for valuable discussions and support. We also thank Katarzyna Agata Wiśniewska and Julia Schleuning for their assistance during sampling for our tracer metabolomics experiments. Finally, we would like to thank and memorialize Palle Serup, who contributed significantly to the first half of the project until his passing.

## Authors contributions

ASPA, CLFH and CH conceived and designed the overall study. ASPA performed experiments and data analysis and interpretation. MBL and CTM generated and supported data analysis of the proteomics data. KHL supported the bulk RNA-seq analysis. ASPA, CLFH, EK, and JJZ conceived and designed the isotope-tracer metabolomic study. TM contributed to the tracer metabolomics analysis. ASPA and EK analyzed and interpreted the stable-isotope tracing data. ASPA and CLFH wrote the manuscript with input from all authors. EK supervised the stable-isotope tracing and metabolomics components. CLFH, CH, PS, and JJZ supervised the project. All authors, PS excluded, reviewed and approved the final manuscript.

## Declaration of generative AI and AI-assisted technologies in the writing process

AI assistance (Claude 4.6 and M365 Copilot) was used only to improve readability and language during manuscript preparation. No AI tools were used to create any of the data or to create or alter images.

## Statistical analysis

All data were collected from at least three independent experiments unless otherwise specified. Data are presented as mean ± standard deviation unless otherwise specified. Statistical comparisons were performed using one-way ANOVA or Welch’s t-test, as specified in the figure legend. Statistical analysis was performed in GraphPad Prism. “N” denotes the number of independent experiments, whereas “n” denotes number of technical replicates.

## Supplementary information

Supplementary Figures 1-7

Proteomic pathway analysis, 6,24,48 hours (Excel)

Transcriptomics pathway analysis, 6,24,48 hours (Excel)

Proteomic differential expression analysis, 6,24,48 hours (Excel)

Transcriptomics differential expression analysis, 6,24,48 hours (Excel)

## Method Text

### Expansion of hPSCs

Human embryonic stem cells, NN1 hESC (Gender: XY, P15), and the induced pluripotent stem cells (hiPSC) SB adult 3.1 (SB3.1, P28) (van de Bunt et al. 2016; Gupta et al. 2018) were used in this study. The NN1 hESCs working cell bank was karyotypically normal (20 cells examined, Cell Guidance Systems). hPSCs were maintained in tissue-culture flasks pre-coated with iMATRIX-511MG (0.5µl/cm^2^, Matrixome #890-005) in iPS-Brew (Miltenyi #170-076-317, #170-076-318) supplemented with TGF-β1(16.8 U/mL, Miltenyi #170-076-166) and Penicillin-Streptomycin (Gibco, 15140-122), referred to as hPSC-medium. hPSCs were cultured at 37°C in a humidified atmosphere with 5% CO₂.

hPSCs were passaged every 3-4 days depending on seeding density. During passages, the hPSC-medium was supplemented with 10 µM Y-27632 ROCKi (Tocris, #1254). For passage, cells were washed in PBS without Ca^2+^/Mg^2+^ (PBS-/-) and enzymatically detached using Accutase (0.02 ml/cm^2^, Stem Cell Technologies, #079200), at 37°C for 4 min. hPSC-medium containing ROCKi was added to neutralize Accutase. Cell suspension was transferred to centrifugation tubes and pelleted at 400g, 5 min. Cell pellet was resuspended in hPSC-medium supplemented with 10µM ROCKi. hPSC viability and cell count were determined using the automatic NucleoCounter® NC200™ or NC-202™ (Chemometec, Denmark). hPSCs were seeded in hPSC-medium supplemented with 10 µM ROCKi into pre-coated flasks, based on the live cell count at a density of 0.012 x10^6^ or 0.022 x10^6^ cells/cm^2^, depending on expansion interval (3-4 days). The medium was changed daily with hPSC-medium without ROCKi.

### Cell cultures

Experiments were initiated from a 4-day passage-interval, as described above. hPSCs were filtered through a 40 µm falcon strainer (Corning, 352340) to ensure a single cell suspension prior to inoculation:

For the 2D culturing hPSCs were seeded at a density of 0.025 x10^6^ cells/cm^2^. The medium was changed daily with hPSC-medium without ROCKi.

For 3D suspension cultures, hPSCs were inoculated with 0.25 x 10^6^ cells/mL into low attachment 6-well plates at a volume of 3 mL per well (0.75 x 10^6^ cells/well). hPSCs were placed on an orbital shaker (CellTron), at a speed of 70 revolutions per minute (RPM). Cells were inoculated in hPSC-medium supplemented with 10 µM ROCKi. The medium was changed after 24 hours with hPSC-medium supplemented with 5 µM ROCKi, unless otherwise stated. When bioreactors were inoculated; single cells were seeded, for DASbox: in 150 ml at an inoculation density of 0.25 x 10^6^ cells/mL unless otherwise stated, medium was changed daily by batch-feeding unless otherwise stated. For DASgib: cells were inoculated in 1 L at a density of 0.5 x 10^6^ cells/mL, with no medium change before sampling 48 hours after inoculation. In experiments where NutriStem medium was used, the hPSC expansion and subsequent culture methods were performed identical to those described for iPS-Brew (hPSC-medium). Samples were collected from each culture system, 2D and 3D suspension, and analyzed using flow cytometry, proteomics, bulk RNA sequencing, SDS-PAGE and western blotting, and isotope-tracer metabolomics.

### Morphology and aggregate size

Brightfield images were captured at different time points (6, 24 and 48 hours) depending on the experiment. Aggregate sizes (only 3D suspension) were monitored using a BIOREP automatic Islet counter (ICC4-115/230). For each analysis 2 technical replicates were analyzed.

### Flow cytometry

For flow cytometry, 2D cultured hPSCs were dissociated as described for cell passaging to obtain a single cell suspension and filtered through a 40 µm falcon strainer. For 3D suspension, aggregates were pooled from a few wells in a falcon tube. Wells were washed with PBS-/- and aggregates were sedimented by centrifugation at 100 x g for 1 min. Supernatant was removed; aggregates were washed once in PBS-/- followed by dissociation in 2 ml Accutase at 37℃ on the orbital shaker for 4 min. Aggregates were carefully pipetted to obtain a single cell suspension, and hPSC-medium was used for neutralization. The suspension was filtered through a 40 µm falcon strainer to ensure a single cell suspension.

For both 2D and 3D suspension, cells were counted before continuing. The cell counts were used to evaluate cell growth in both systems.

For flow cytometry approximately 2.5 x 10^6^ cells were stained with a live/dead dye (NEAR-IR, Life Technologies, #L34976) before fixation in 1 ml 4% formaldehyde (Cell Signaling, #47746P) for 20 min. After fixation samples were washed in 12 ml PBS-/- with 1% BSA (Wash buffer), centrifuged at 800 x g for 3 min and resuspended in wash buffer at a density of 2.5 x 10^6^ cells/ml.

2.5 x 10^5^ fixed cells/stain were permeabilized using PBS-/- with 0.2% Triton-X (Thermo Scientific™, #85111) and 2% BSA (Sigma-Aldrich, #A7284) (Perm buffer) for 30 min at room temperature (RT) in the dark. Cells were subsequently incubated with OCT3/4-AF647 (BD Biosciences, #560307) and SOX2-V450 (BD Biosciences, #561610) for 30 min, RT, in the dark. Antibodies were diluted 1:25 and 1:80 respectively. Hereafter, samples were washed three times using wash buffer and filtered through a 40 µm filter plate before acquisition using CytoFlex Flow Cytometer (Beckman Coulter, California, USA). Samples were analyzed within 24 hours following staining. For each analysis, one unstained control and relevant fluorescence-minus-one (FMO) controls were included. Compensation beads (Invitrogen^TM^ #01-3333-42 or #A10346) were always included by staining 1 drop for 15-30 min with each antibody followed by washing and acquisition. Data analysis was performed using FlowJo (version 10.8.1).

### Proteomics

For proteomics, 2D hPSCs were washed three times with ice-cold PBS -/- and harvested using cell-scrapers in 5 ml PBS-/- before transferring to a Falcon tube. The culture flask was washed with an additional 5 ml PBS-/-. Cells were pelleted by centrifugation 400 x g for 4 min. Supernatant was discarded, and the cell pellet was resuspended in 750 µl ice-cold PBS-/- and transferred to low-bind protein Eppendorf tubes. For the 3D suspension cultured hPSCs, aggregates were transferred to a falcon tube. The 6 well plates were washed in ice-cold PBS-/-. Aggregates were sedimented by centrifugation 100 x g for 1 min and supernatant was removed. The wells and aggregates were washed in 5 ml PBS-/- twice. Aggregates were centrifuged 400 x g for 5 min and resuspended in 750 µl ice-cold PBS-/- and transferred to low-bind protein Eppendorf tubes. Samples were centrifuged at 800 x g for 5 min. The supernatant was discarded and cell pellet was snap-frozen and stored at -80℃.

Cell pellets were homogenized in 500 µl lysis buffer (6M Guanidine-HCl, 5 mM Tris(2-carboxyethyl)phosphine hydrochloride (TCEP, Thermo Scientific, #77720), 10 mM 2-Chloroacetamide (CAA, Aldrich, #C0267), 100mM Triethylammonium bicarbonate (TEAB, Sigma-Aldrich, #90360) at 90°C. Lysates were further disrupted by sonication (Cup horn sonicator, 3×30s, 30s cool down). Protein concentration was determined by NanoDrop A280 absorbance. A total of 150 µg protein was diluted with 100mM TEAB followed by digestion at 37°C for 16 hours using Trypsin/Lys-C (Trypsin/LysC mix, MS grade, Promega, #V5073) in a 1:50 ratio. Digestions were stopped by adding Trifluoroacetic Acid (TFA, Fisher, #A116) to a final concentration of 1%. Peptides were subsequently TMT-labelled. Peptides were cleaned using 50 mG tC18 SepPak columns (Waters, #WAT054960) following vendor protocol. Eluted peptides were dried and resuspended in 50 mM TEAB. 20 µg peptide from each sample were labelled in groups of 10 using TMT10plex (Thermo Scientific, #90110) followed by a quench with hydroxylamine. Each group was then pooled and cleaned on SepPak and dried. Samples were fractionated via high-pH offline fractionation. Dried peptides were resuspended in 50 µl HpH buffer A (5mM Ammonium bicarbonate pH 8 (ABC; Sigma Aldrich #A6141)) and subjected to reverse-phase high-performance liquid chromatography on a Dionex UltiMate 3000 HPLC system (Thermo Scientific, USA) using a Waters CSH C18 analytical column (1 mm x 150 mm, 130 Å, 1.7 µm; #186005294). Peptides were loaded onto the column in Buffer A and eluted at 30 µL/min in a gradient from 5% to 25% Buffer B (5 mM Ammonium bicarbonate pH 8 (Sigma-Aldrich, #A6141) (in 90% acetonitrile (ACN, Fisher, #A955-212) over 55 min. For each group 23 fractions were collected and speedvaced to near dryness. Fractions were reconstituted in 7 µl 2% ACN, 0.1% FA and 5 µl of the peptide eluate were separated on a 60 min gradient on an EASY-nLC 1200 (Thermo Fisher) using an Evosep 15 cm column (EvoSep Performance #EV-1137). MS analysis was performed on an Exploris 480 mass spectrometer (Thermo Fisher) using Data-Dependent Acquisition method with MS1 scan settings of 350-1400 m/z mass range, resolution 60000, and an AGC setting of 5e6. MS2 was obtained at resolution 30000, fixed injection time of 54 ms, AGC 5e5 with a normalized collision energy of 33.

Raw data were searched against the human UniProt/SwissProt database using MaxQuant v1.6.1 with the Andromeda search engine with FDR of 0.01 for protein and peptides. The MS/MS spectra were searched with variable modifications Oxidation (M), acetyl (N-term), fixed modifications matching TMT10plex and Carbamidomethyl (C) and a specific Trypsin/P pattern with a maximum of two missed cleavages. Peptide length was limited to a minimum of 7 amino acids. Further data analysis was performed using Perseus (Version 1.6.15.0) together with annotated metadata. All data were Log_2_-transformed and proteins identified by site only, site, reverse and potential contaminants were removed. Using the Swiss-Port database, we added Gene Ontology (GO) annotations, biological process, cellular compartments, and molecular function, together with pathways and keywords. Only proteins with valid quantitative values in every sample were retained for analysis.. Data were further normalized using Quantile normalization to reduce within-batch effect, followed by ComBat correction for between-batch effects. Differential expression analysis was performed with an FDR threshold of 0.05. Visualization and pathway enrichment analysis were performed in R 4.2.0.

### Bulk RNA-sequencing

For bulk RNA sequencing, cell cultures were dissociated as described to obtain a single cell suspension. For each condition, 2 x 10^6^ cells were transferred to a falcon tube and centrifuged 400 x g for 5 min. Supernatant was discarded, and the cell pellet was resuspended in 2 ml Stem CellBanker (AMS BIOTECHNOLOGY (EUROPE) LIMITED, 11890) for cryopreservation in vial containing 1 x 10^6^ cells /tube. Cells were stored in liquid nitrogen until further processing.

All samples were thawed and lysed using RLT buffer (QIAGEN, #79216) and RNA was extracted using RNeasy Plus Micro Kit (QIAGEN, #74034) followed by determination of RNA concentration and quality using Nanodrop and Bioanalyzer (Agilent). RNA integrity numbers (RINs) were 10 for all assessed samples, except for three samples with unavailable RIN values. However, result inspection did not indicate reduced RNA quality. RNA was shipped for sequencing at Eurofins Genomic using Genome Sequencer Illumina NovaSeq 600 S4 PE150 XP, and standard-specific cDNA library. Sequencing depth was >33,000,000 reads/sample.

Data analysis was performed using R 4.2.0 and the DESeq2 package (Love et al., 2014). Data was normalized and a study design generated based on assessment of the differences found from a principal component analysis (PCA). Gene set enrichment analysis (GSEA) using KEGG pathways was performed at all time points comparing 2D and 3D cultured hPSCs. For visualization the following packages were used: tidyverse (Wickham et al., 2019), circlize (Gu et al., 2014), ComplexHeatmap (Gu et al., 2016) EnhancedVolcano (Blighe et al., 2025), clusterProfiler (Xu et al., 2024), org.Hs.eg.db (Carlson, 2025), enrichplot (Yu, 2026), and Cairo (Urbanek and Horner, 2025).

### Western blotting

hPSCs for SDS-PAGE/Western blot analysis were harvested as described for proteomics. Cell pellets were lysed using RIPA buffer (Thermo Scientific™, #89900) containing phosphatase (Roche, 04 906 837 001) and protease (Roche, #04 693 124 001) inhibitors. Protein concentrations were determined using the Pierce™ BCA Protein Assay Kit (Thermo Scientific™, #23227). 10 µg protein/sample were separated by SDS-PAGE and transferred to PVDF membranes using the iblot PVDF mini stacks (Invitrogen^TM^, #IB24002). Blocking and dilution of antibodies were performed in Pierce Fast Blocking Buffer (Invitrogen^TM^, #37575). Membranes were stained with primary target antibody overnight at 4℃, washed three times, followed by incubation with an HRP-conjugated secondary antibody for 1 hour at RT. For the loading control membranes were incubated 1hr for both primary and secondary antibodies. Super signal Kit west Dura extended duration substrate (Invitrogen^TM^, #34075) was added to the PVDF membranes and signal was detected using the Amerhams Imager 600. Signal intensity was normalized to the loading control (HSP90) using ImageJ (version 1.54d).

### Lactate and Glucose measurements

Yield coefficient of Lactate from Glucose Y(qLac/qGlc) was determined as described in (Kropp *et al*., 2016; Manstein *et al*., 2021). Briefly, cell culture medium was analyzed for glucose and lactate concentrations using a RAPIDPoint® 500e Blood Gas System (Siemens). Medium was collected prior to inoculation or medium changes as reference samples. Conditioned medium was analyzed 24 and 48 hours following inoculation. At the same time a cell yield assessment was performed to allow normalization.

The mean cell concentration, *x̅*_*t*__*n*+1_ [cells/L], was calculated using the following formula

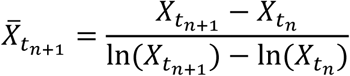

X represents the cell concentration [cells/L].

The specific glucose consumption rate (qGlc) [mmol cell^-1^ day^-1^] for the sampling day was calculated as

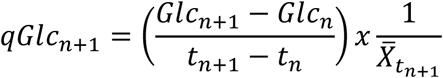

Glc represents the glucose concentration [mmol/L], t represents the process timepoint [day]. The specific Lactate production rate (qLac) [mmol cell^-1^ day^-1^] for the sampling day was calculated as

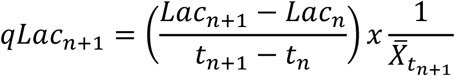

Lac represents lactate concentration [mmol/L].

The yield coefficient of Lactate from Glucose Y(qLac/qGlc) [-] was calculated as

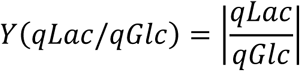

### Stable isotope metabolomics

5 hours prior to harvest, medium was changed to unlabeled or isotope-labeled DMEM/F12 (Ref#06-1170-94-1A), 20 mM Glucose (Unlabeled: GIBCO, #A2494001, labeled: ChemSupport, CLM-1396-1 or 1-978-749-8000) and 4 mM glutamine (Unlabeled: ThermoFisher Scientific, #35050038, labeled: ChemSupport, CLM-1822-H-0.1mg, 18411-19-1) final concentration, iPS-Brew GMP Supplement R (50x dilution) and TGF-β1 (16.8 U/mL). After 5 hours, samples were washed three times with PBS. Leftover PBS was removed followed by addition of 1 ml ice-cold 90% Methanol. For unlabeled samples methanol was supplemented with isotopically labeled internal controls. 2D cultured cells were collected by scraping. 3D cultured aggregates were transferred to a 40 µm cell strainer to retain aggregates and remove cell culture medium. The cell strainer was moved to a clean 6 well plate. A volume of 0.5 ml ice-cold 90% Methanol was added to the filter with aggregates and the filter was scraped. An extra 0.5 ml of ice-cold 90% Methanol was added before collection. Methanol samples were collected in Eppendorf tubes, and snap-frozen and stored at –80℃ until intracellular metabolite extraction. For each culture system a sample was harvested for protein extraction and quantification using Pierce™ BCA Protein Assay Kit.

Samples were thawed on ice, and snap-frozen in liquid nitrogen, thawed and vortexed. This cycle was repeated three times followed by incubation on ice for 1hr, prior to centrifugation at 15000 rpm for 15 minutes at 4℃. Supernatant was collected in fresh tubes and stored until further processing.

TCA metabolites were analyzed according to Kafkia et al. (Kafkia *et al*., 2026). The LC-MS analyses were performed using a 1290 Infinity III Bio ultra-high-performance liquid chromatography (UHPLC) system (Agilent Technologies, Waldbronn, Germany) coupled to a timsTOF Pro 2 mass spectrometer (Bruker Daltonics, Bremen, Germany). Central carbon metabolites were derivatized with 3-nitrophenylhydrazine (3-NPH) according to Hodek et al. (Hodek et al., 2023), with minor modifications. Briefly, 20 µl of 120 mmol L-1 EDC (dissolved in 6% pyridine and 50% methanol) and 20 µl of 200 mmol L-1 3-NPH (dissolved in 50% methanol) were consecutively added to 20 µl of reconstituted samples in 1:1 methanol:water (v/v). The sample was incubated at RT (21℃) for 60 minutes, and afterwards 40 µl of 0.05 mg mL^-1^ BHT (dissolved in 30% pure methanol) were added to the samples and vortexed. 2 µl of each sample were analyzed on an Acquity UPLC HSS-T3 column (100 ×2.1 mm, 1.8 μm, Waters, MA, USA) by using gradient elution of 0.1% formic acid (v/v) in water as mobile phase A and 0.1% formic acid in acetonitrile as mobile phase B. The flow rate was set at 0.35 mL min^-1^ with the following gradient: 0 min (5% B), 12 min (100% B), 13 min (100% B), 14 min (5% B), 17 min (5% B). Column and autosampler were kept at 30℃ and 4℃, respectively. The mass spectrometer was operated in negative ionization mode. Ions were generated with a vacuum insulated probe heated-electrospray ionisation (VIP-HESI) source. Detection of the mass-to-charge ratio (m/z) of ions was set from 50 to 1000, and acquisition rates were set to 2 Hz, and the resolution was approximately 60,000. For mass calibration 50 µl internal calibrant of 10 mM Na-formate was injected at the beginning of each analysis. Data acquisition was performed with Control version 6.0 and Bruker Compass HyStar version 5.0 (Bruker Daltonics, Bremen, Germany). Data processing was performed with Bruker TASQ 2025b.

Total carbon contribution was calculated using the following equation (Nanchen et al., 2007):

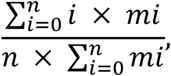

where *n* denotes the total number of carbons in the metabolite, *i* denotes the different mass isotopologues and *m* corresponds to the abundance of a certain mass.

