## Supplementary Figures 1-7 for "Suspension culture drives metabolic rewiring in human pluripotent stem cells"

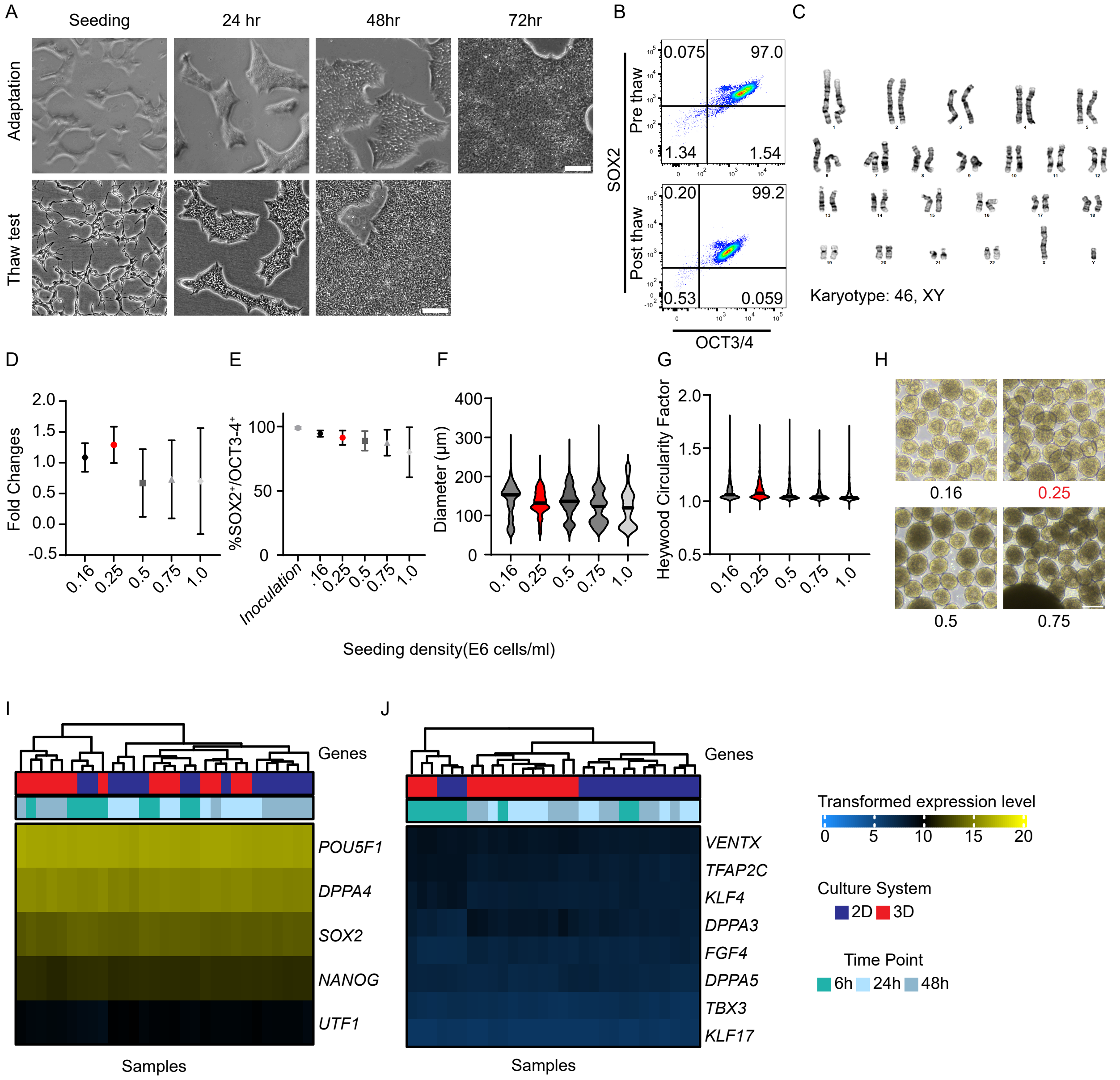


**Supplementary Figure 1. Characterization of human embryonic stem cell working cell bank and suspension culture conditions.**

A–C) Human embryonic stem cells (hESC) were adapted to the medium, GMP iPS-Brew, prior to cryopreservation as the working cell bank. A) Representative bright field images at seeding, 24, 48, and 72 hours during media adaptation (between passages 2 and 3) and following thaw of the working cell bank. Scale bar: 200 µm. B) Flow cytometry density plots of SOX2 and OCT3/4 before cryopreservation and following thaw. C) Karyotype analysis (G-banding) of the working cell bank.

D–H) Five seeding densities (0.16, 0.25, 0.5, 0.75, and 1.0 × $10^6$ cells/mL) were evaluated for 3D suspension culture. The selected condition (0.25 × $10^6$ cells/mL) is highlighted in red throughout panels. D) Fold change in cell count between inoculation and 48 hours. E) Pluripotency assessed by flow cytometry based on co-expression of SOX2 and OCT3/4. F) Aggregate diameter measured using the BIOREP islet counter. G) Aggregate circularity assessed using the Heywood circularity factor. H) Representative bright field images of aggregates at seeding densities of 0.16, 0.25, 0.5, and 0.75 × $10^6$ cells/mL. Scale bar: 200 µm. N = 2–3. I-J) Bulk RNA sequencing analysis of primed and naïve pluripotency markers shown as heatmaps of transformed expression levels, categorized by culture system and time point. I) Primed pluripotency, J) Naïve pluripotency. N=5

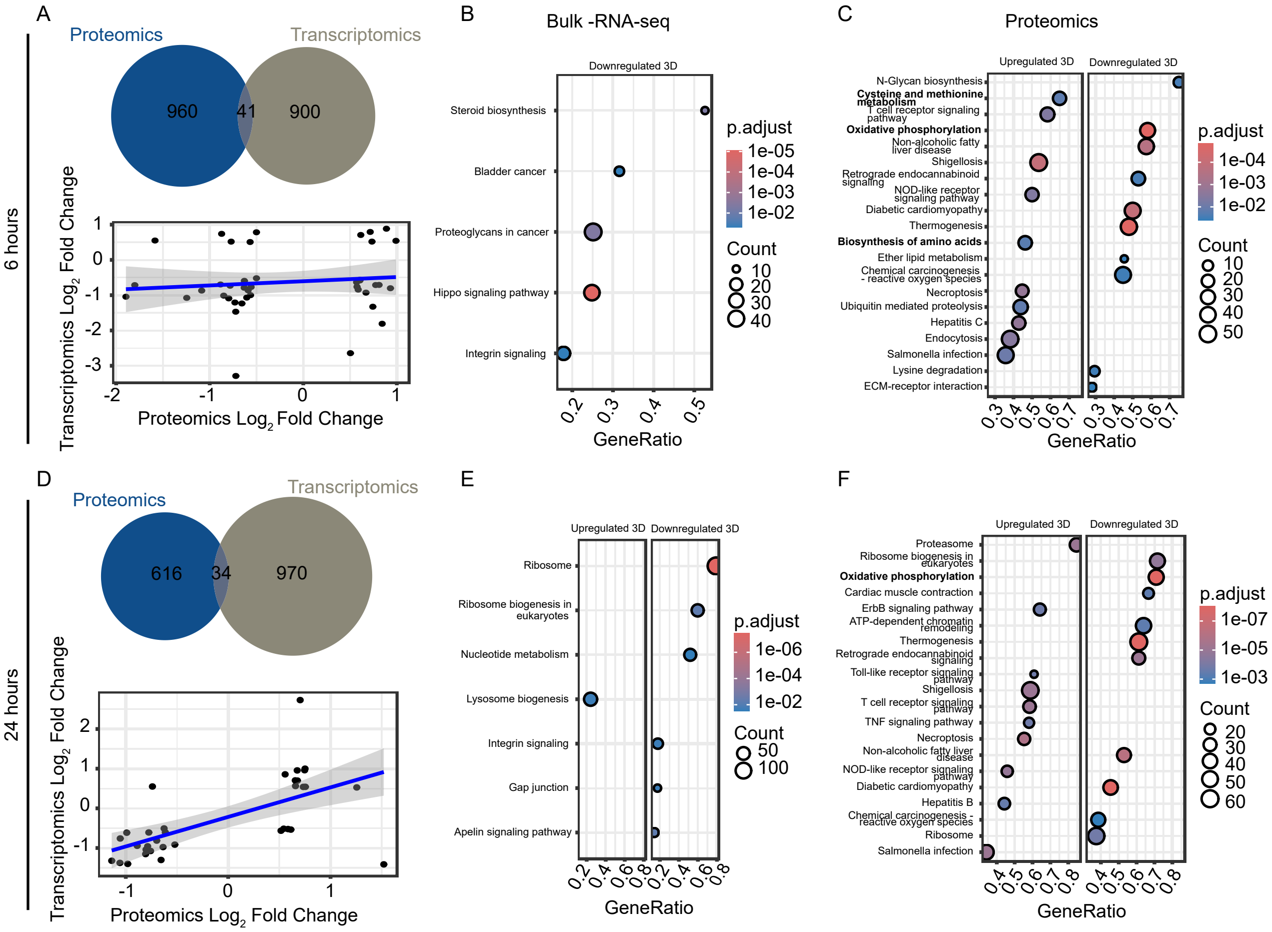


**Supplementary Figure 2. Correlation between proteomics and transcriptomics and KEGG pathway enrichment analysis at 6 h and 24 h post-inoculation.**

A–C) Analysis at 6 hours and D-F) 24 hours post-inoculation. A and D) Venn diagrams (top) illustrate the overlap between differentially expressed proteins and genes. Correlation plot (bottom) of Log2 fold changes between proteomics and transcriptomics for overlapping hits. The blue line represents a linear regression fit. B and E) KEGG pathway enrichment analysis of transcriptomic data showing the regulated pathways. C and F) KEGG pathway enrichment analysis of proteomic data, showing the top regulated pathways in 3D suspension compared to 2D culture. Dot size reflects gene ratio, and color indicates adjusted p-value.

N = 5 biological replicates. See also Figure 2.

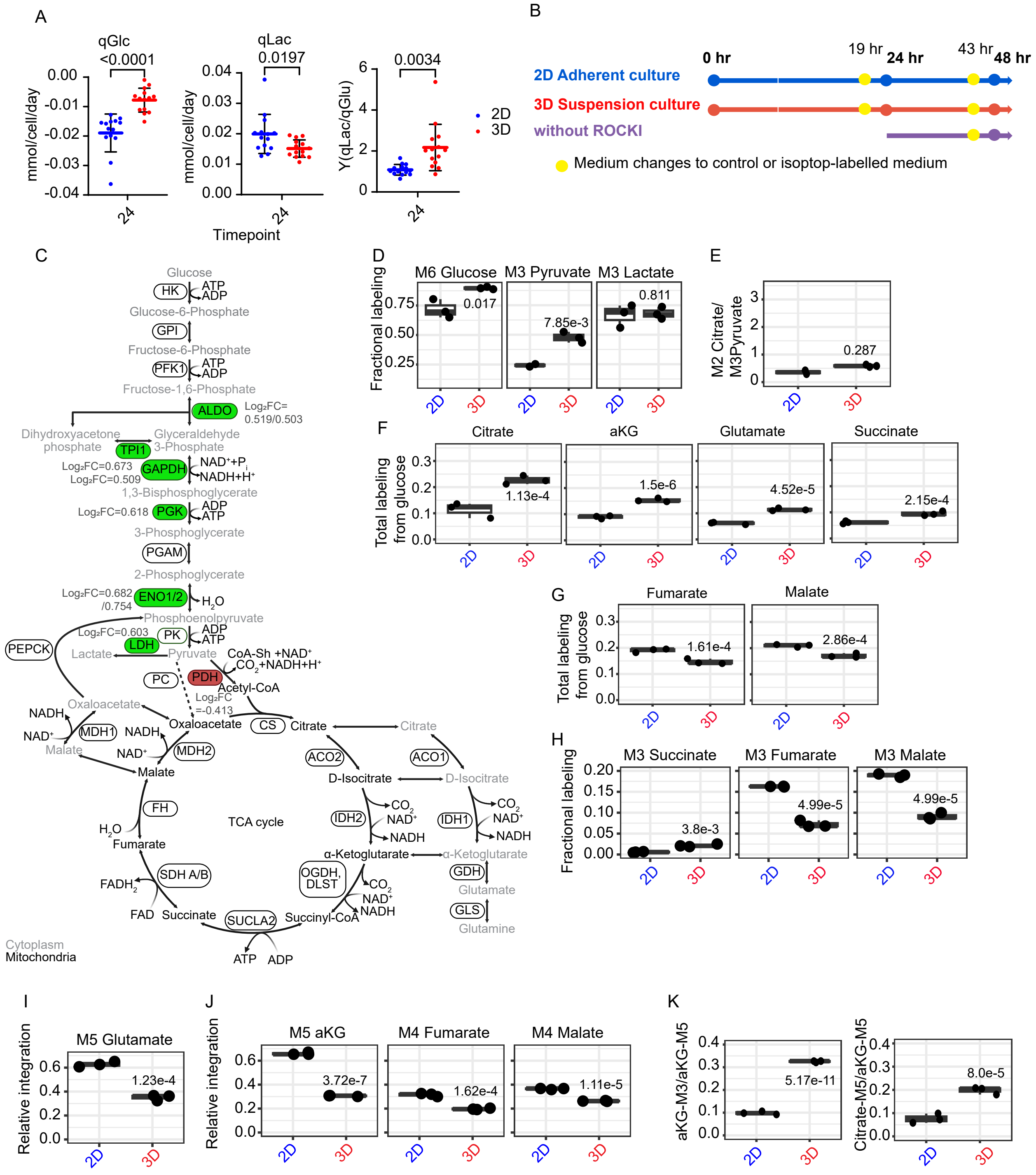


Supplementary Figure 3. The legend is provided on the next page.

**Supplementary Figure 3. Increased glycolytic and metabolic activity in 3D cultured hPSCs observed at 24 hours.**

A) Spent-medium analysis of glucose consumption (qGlc), lactate production (qLac), and lactate yield from glucose (Y(qLac/qGlc)) at 24 hours. N = 14; data presented as mean ± SD with individual biological replicates. Statistical analysis by Welch's t-test.
B) Schematic illustration of the experimental design for stable-isotope tracer metabolomics.
C) Schematic illustration of glycolysis and the TCA cycle highlighting enzymes with increased (green) and reduced (red) abundance in 3D suspension culture at 24 hours.

D-H) [U-$^{13}$C]-glucose tracing. D) Fractional labeling of M6 Glucose, M3 Pyruvate and M3 Lactate, E) ratio of M2 Citrate to M3 pyruvate. F and G) Total labeling of Citrate, α-KG, glutamate, and succinate, Fumarate and Malate from glucose. H) Fractional labeling of M3 Succinate, M3 Fumarate, and M3 Malate.
I-K) [U-$^{13}$C]-glutamine tracing. I) Fractional labeling of M5 glutamate. J) Fractional labeling of M5 α-KG, M4 fumarate, and M4 malate reflecting glutamine entry into the TCA cycle. K) Ratio of M3 α-KG to M5 α-KG and the ratio of M5 citrate to M5 α-KG.
n = 3 technical replicates; data presented as mean with individual replicates. Statistical significance was assessed using one-way ANOVA. P-values are indicated for each comparison.

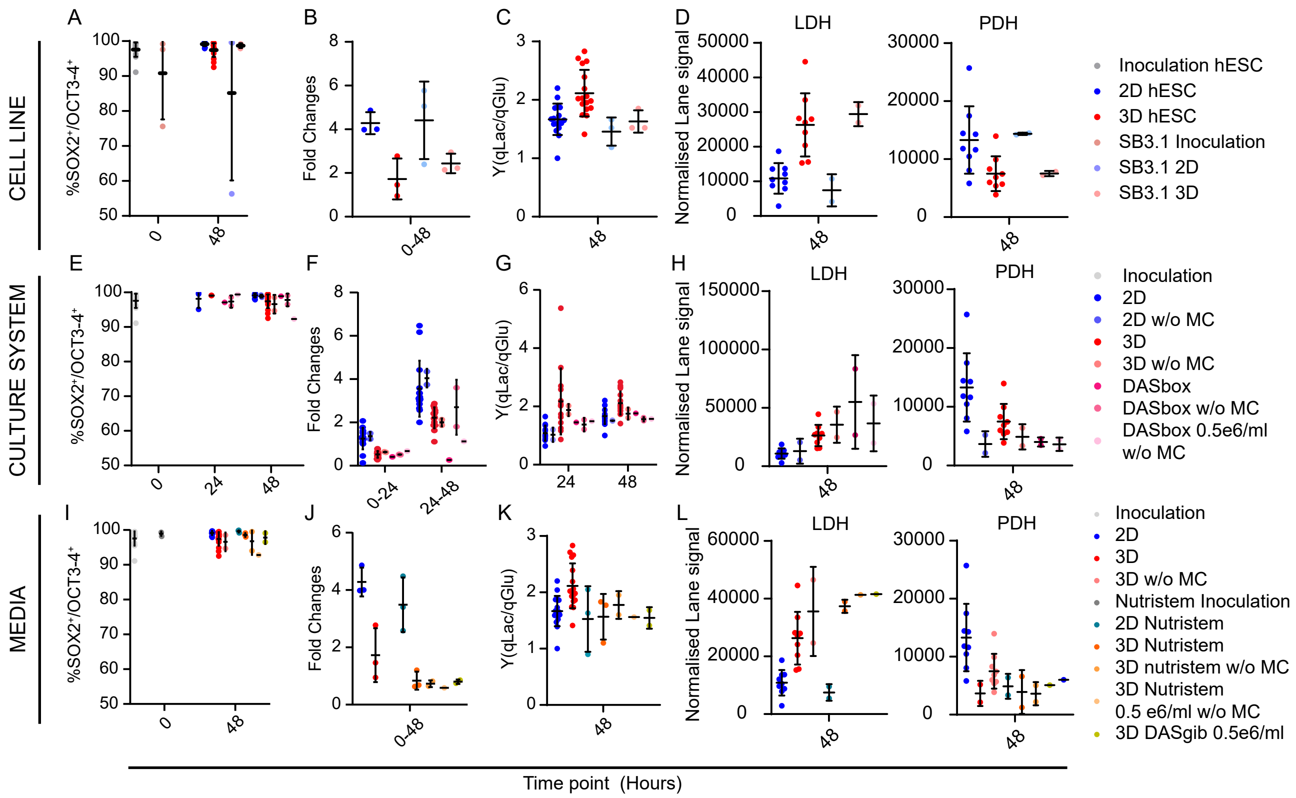


**Supplementary Figure 4. Metabolic changes are observed across cell lines, culture systems, and media formulations.**

A–D) To evaluate whether the observed metabolic changes were independent of cell line, the iPSC line Adult SB3.1 was evaluated alongside the hESC line used in this study, after 48 hours in 2D and 3D suspension culture.

E–H) Results were obtained in a small-scale 6-well plate suspension culture system, we evaluated if our findings translated to a stirred tank bioreactor system (DASbox). Here we have included conditions with or without medium changes (w/o MC), because process parameters often include omitting the 24-hour medium changes in STB.

I–L) To assess dependence on medium and culture system, experiments were repeated using Nutristem medium and the STB system, DASgib.

A, E, I) Pluripotency assessed by co-expression of SOX2 and OCT3/4 by flow cytometry. B, F, J) Cell growth expressed as fold change. C, G, K) Yield specific lactate-to-glucose ratio [Y(qLac/qGlc)]. D, H, L) LDH and PDH protein abundance quantified by western blotting (normalized lane signal) at 48 hours.

Data are presented as mean ± SD. N = 1–3 biological replicates depending on the measurement

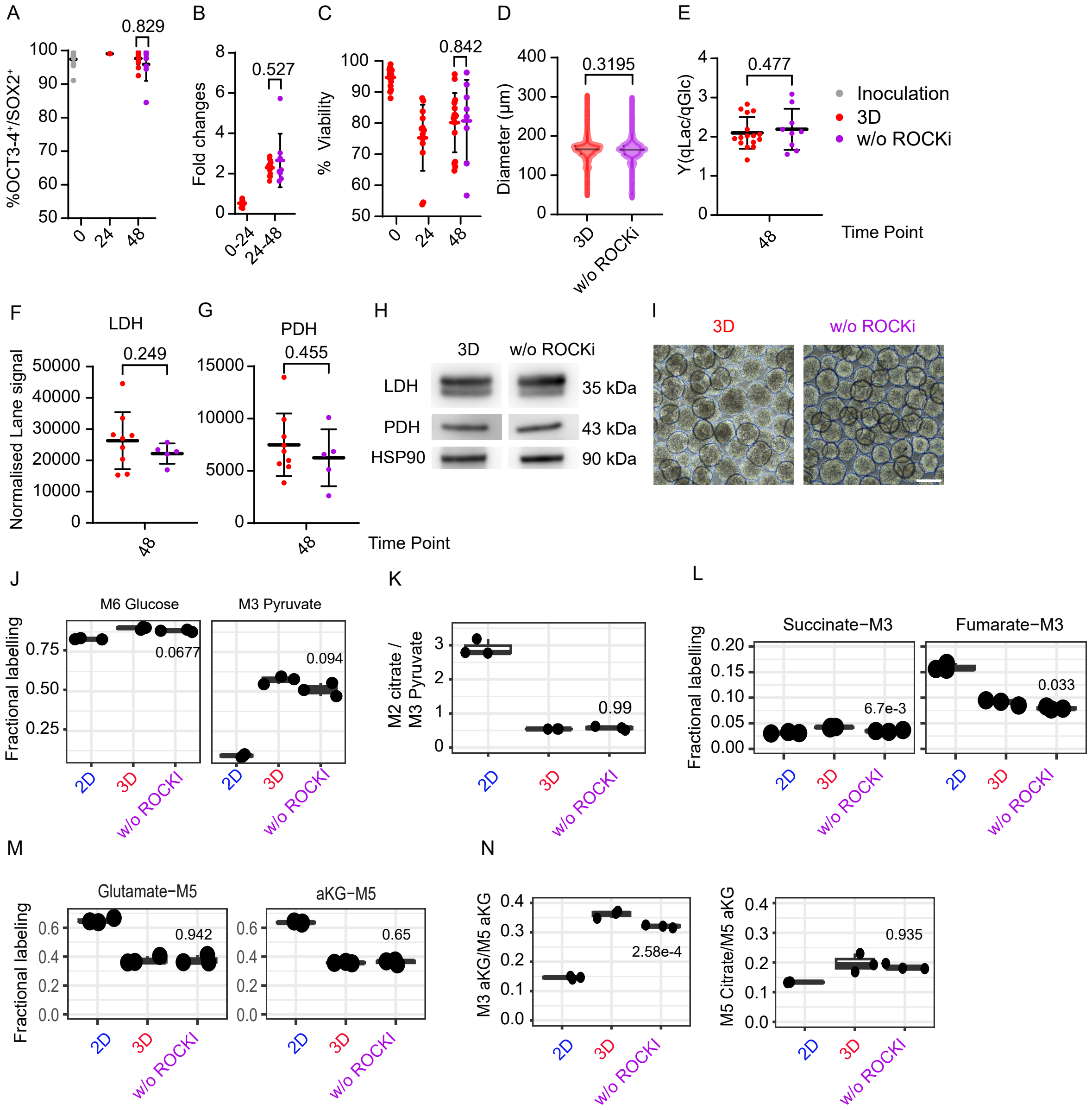


**Supplementary Figure 5. The legend is provided on the next page.**

**Supplementary Figure 5. Limited effect of ROCK inhibition on the metabolic profile of hPSCs in suspension culture.**

5µM ROCKi (Y-27632) was added to 3D suspension cultures from 24 to 48 hours. Basal culture parameters were assessed comparing 3D with and without ROCKi: A) Pluripotency assessed by flow cytometry for co-expression of SOX2 and OCT3/4. B) Fold change in cell count. C) Cell viability. D) Aggregate size measured using the BIOREP islet counter. N = 17 (3D) and N = 10 (3D without ROCKi).

E) Spent-medium analysis quantifying lactate yield from glucose Y(qLac/qGlc). F) LDH and G) PDH protein abundance quantified by western blotting. H) Representative western blots of LDH, PDH, and HSP90 (loading control). N = 9 (3D) and N = 5 (3D without ROCKi).

I) Representative brightfield images at 48 hours of 3D cultures with and without ROCKi. Scale bar: 200 µm.

Data presented as mean ± SD with individual replicates. Statistical analysis by Welch's t-test.

J-L) [U-$^{13}$C]-glucose tracing. J) Fractional labeling of M6-glucose and M3 pyruvate. K) Ratio of M2 Citrate to M3 pyruvate. L) Fractional labeling of M2 succinate and M3 fumarate.

M-N) [U-$^{13}$C]-glutamine tracing. M) Fractional labeling of M5 glutamate and M5 α-KG. N) Ratio of M3 α-KG and M5 citrate to M5 α-KG.

n = 3 technical replicates; data presented as mean with individual replicates. Statistical significance was assessed using one-way ANOVA, p-values for the comparison between 3D with and without (w/o) ROCKi are shown in each plot.

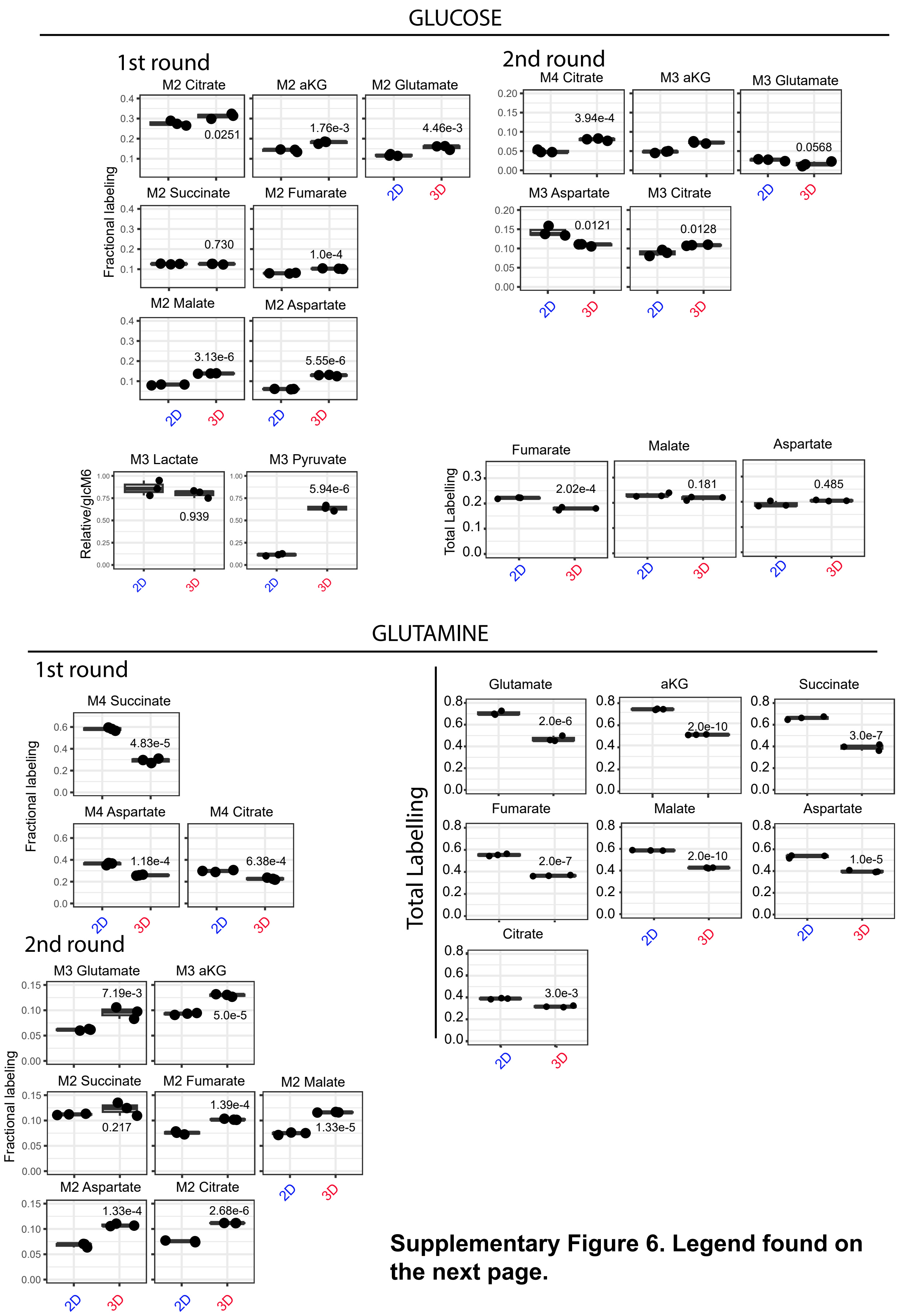


**Supplementary Figure 6. Legend found on the next page.**

**Supplementary Figure 6. Glucose- and glutamine-derived carbon incorporation into TCA metabolites in 2D and 3D hPSC cultures after 48 hours.**

Fractional isotopologue labelling and total carbon labelling of intracellular metabolites following tracing with [U-$^{13}$C]-glucose or [U-$^{13}$C]-glutamine in hPSCs cultured under 2D adherent and 3D suspension conditions.

For glucose tracing, M2 isotopologues indicate first-round incorporation of glucose-derived carbon into TCA cycle intermediates, whereas M3 and M4 isotopologues reflect the second round of labelling through the TCA cycle. Total labelling of fumarate, malate, and aspartate is shown below. For glutamine tracing, M4 and M5 isotopologues represent first-round incorporation of glutamine-derived carbon into TCA cycle-related metabolites, while M2 and M3 isotopologues indicate further TCA cycling and downstream carbon redistribution. Total labelling of glutamate, α-ketoglutarate (aKG), succinate, fumarate, malate, aspartate, and citrate is shown on the right.

n = 3 technical replicates; data presented as mean with individual replicates. Statistical significance was assessed using one-way ANOVA, p-values are shown in each plot for the comparison of 2D to 3D suspension cultures.

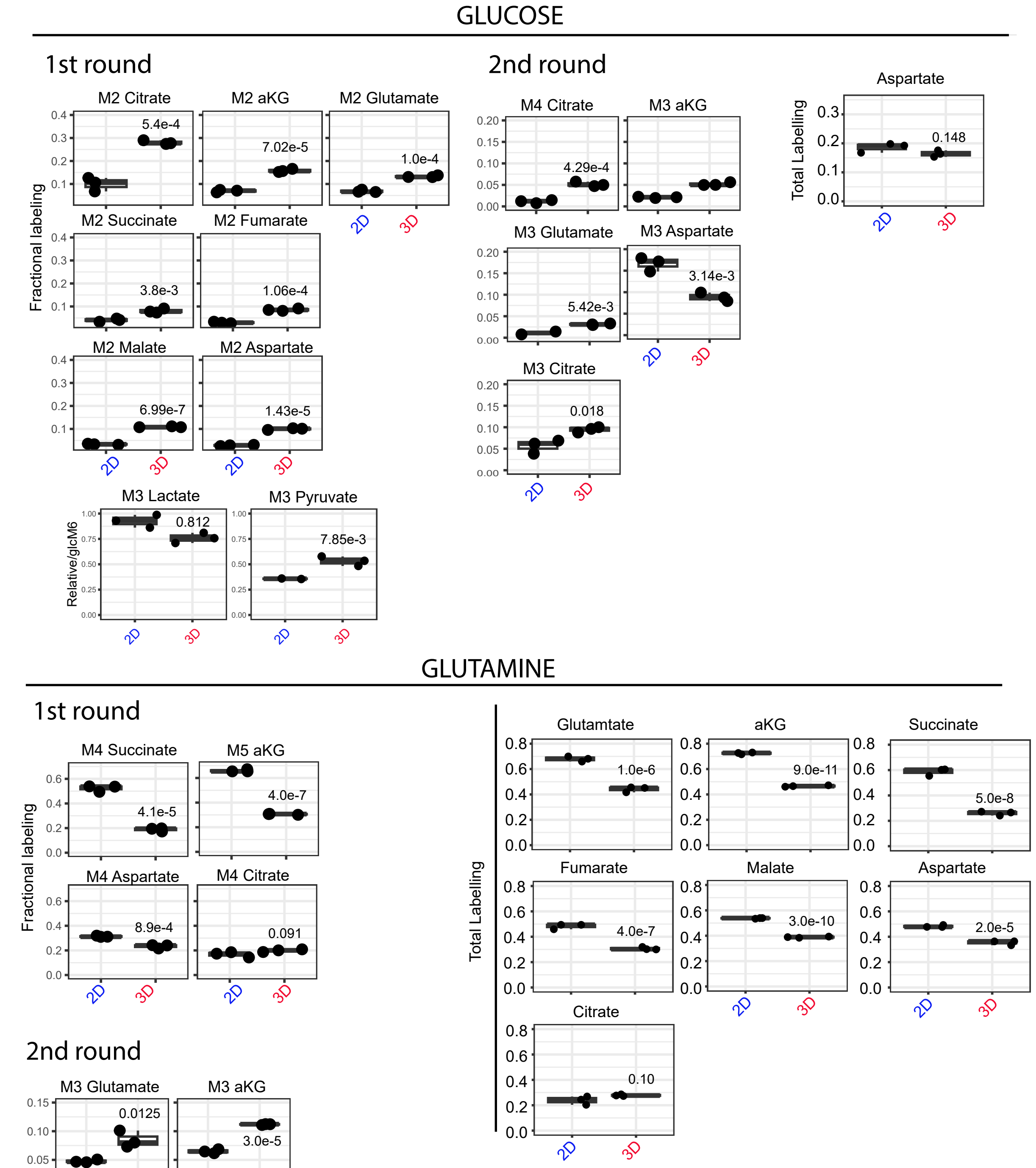

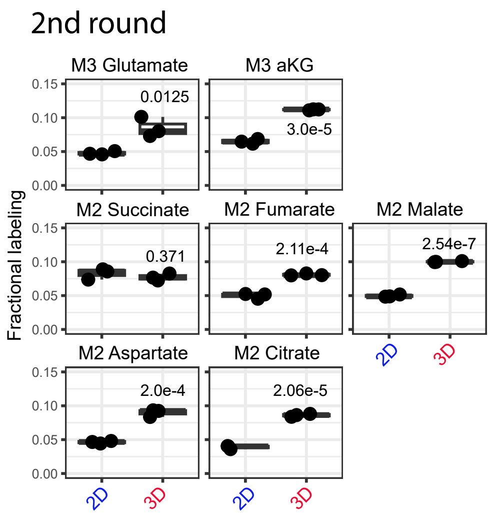


**Supplementary Figure 7. The legend is provided on the next page.**

**Supplementary Figure 7. Glucose- and glutamine-derived carbon incorporation into TCA cycle intermediates in 2D and 3D hPSC cultures at 24 hours.**
Fractional isotopologue labelling and total carbon labelling of intracellular metabolites with [U-$^{13}$C]-glucose or [U-$^{13}$C]-glutamine in hPSCs cultured under 2D adherent and 3D suspension conditions.
For glucose tracing, M2 isotopologues represent first-round incorporation of glucose-derived carbon into TCA cycle intermediates, whereas M3 and M4 isotopologues reflect further the second round TCA cycle and carbon redistribution. Total labelling of aspartate is shown on the right. The ratios of M3 lactate and M3 pyruvate to M6 glucose are also shown.
For glutamine tracing, M4 and M5 isotopologues represent first-round incorporation of glutamine-derived carbon into TCA cycle intermediates, while M2 and M3 isotopologues indicate second round TCA cycle. Total labelling of glutamate, α-ketoglutarate (aKG), succinate, fumarate, malate, aspartate, and citrate are shown on the right.
n = 3 technical replicates; data presented as mean with individual replicates. Statistical significance was assessed using one-way ANOVA, p-values are shown in each plot for the comparison of 2D to 3D suspension cultures.
